# Exploring the upper pH limits of nitrite oxidation: diversity, ecophysiology, and adaptive traits of haloalkalitolerant *Nitrospira*

**DOI:** 10.1101/2020.03.05.977850

**Authors:** Anne Daebeler, Katharina Kitzinger, Hanna Koch, Craig W. Herbold, Michaela Steinfeder, Jasmin Schwarz, Thomas Zechmeister, Søren M. Karst, Mads Albertsen, Per H. Nielsen, Michael Wagner, Holger Daims

## Abstract

Nitrite-oxidizing bacteria of the genus *Nitrospira* are key players of the biogeochemical nitrogen cycle. However, little is known about their occurrence and survival strategies in extreme pH environments. Here, we report on the discovery of physiologically versatile, haloalkalitolerant *Nitrospira* that drive nitrite oxidation at exceptionally high pH. *Nitrospira* distribution, diversity, and ecophysiology were studied in hypo- and subsaline (1.3-12.8 g salt/l), highly alkaline (pH 8.9-10.3) lakes by amplicon sequencing, metagenomics, and cultivation-based approaches. Surprisingly, not only were *Nitrospira* populations detected, but they were also considerably diverse with presence of members of *Nitrospira* lineages I, II and IV. Furthermore, the ability of *Nitrospira* enrichment cultures to oxidize nitrite at neutral to highly alkaline pH of 10.5 was demonstrated. Metagenomic analysis of a newly enriched *Nitrospira* lineage IV species, “*Candidatus* Nitrospira alkalitolerans”, revealed numerous adaptive features of this organism to its extreme environment. Among them were a sodium-dependent N-type ATPase and NADH:quinone oxidoreductase next to the proton-driven forms usually found in *Nitrospira*. Other functions aid in pH and cation homeostasis and osmotic stress defense. “*Ca.* Nitrospira alkalitolerans” also possesses group 2a and 3b [NiFe] hydrogenases, suggesting it can use hydrogen as alternative energy source. These results reveal how *Nitrospira* cope with strongly fluctuating pH and salinity conditions and expand our knowledge of nitrogen cycling in extreme habitats.

## Introduction

Chemolithoautotrophic nitrite-oxidizing bacteria (NOB) are key players of the nitrogen cycle in virtually all oxic habitats including soil, freshwater and marine ecosystems, engineered environments, and geothermal springs [1–9]. By catalyzing the second step of nitrification, NOB are the main biological source of nitrate, which is an important source of nitrogen and a terminal electron acceptor used by a plethora of other organisms. In most terrestrial and engineered environments, the predominant known NOB are uncultured members of the genus *Nitrospira* [1, 10–13]. Within this highly diverse genus, six phylogenetic lineages (named lineage I to VI) have been described, some of which seem to colonize distinct habitat types [1, 4, 8, 14, 15]. Recent studies revealed an unexpected metabolic versatility of *Nitrospira* beyond nitrite oxidation, such as aerobic growth on hydrogen or formate [16, 17] and, most surprisingly, the capability of complete ammonia oxidation to nitrate by some representatives (the comammox organisms) [18, 19].

Haloalkaline systems are highly productive environments that harbor diverse, haloalkaliphilic microbial communities capable of rapid biogeochemical cycling [20–28], but knowledge of the responsible microbes and their ecology, in particular of NOB, is fragmentary [21, 27, 29, 30]. In a pioneering study, the hitherto only known facultatively alkaliphilic nitrite oxidizer, *Nitrobacter alkalicus*, was isolated and analyzed regarding its morphology and tolerance towards elevated pH of around 10 [29].

Shallow, saline-alkaline lakes are a characteristic of the Pannonian steppe in Central Europe – an ecosystem which extends into eastern Austria and is protected in the national park “Neusiedler See - Seewinkel”. The salinity of these lakes varies within the hyposaline range and the pH is generally above 9 [31, 32]. These lakes exhibit a high turbidity caused by inorganic suspended particles and/or high humic substance content and frequently dry out during summer months [31]. Plant material of the shoreline vegetation and excrement of aquatic birds provide organic carbon and inorganic nitrogen and phosphorous inputs [33, 34]. Taken together, shallowness, intermittent character (periodic desiccation), high turbidity, alkaline pH, polyhumic organic carbon concentration, hypertrophic conditions and during summer high daily water temperature fluctuation create multiple extreme environmental conditions in these lakes [35].

In the present study, we obtained deeper insights into the biology of *Nitrospira* in haloalkaline systems. An investigation of the NOB community structure in sediments of saline-alkaline lakes in the national park “Neusiedler See – Seewinkel”, Burgenland, Austria (Fig. 1), by amplicon sequencing subsequently allowed for the targeted study of the ecophysiology and genomic adaptations in newly discovered alkalitolerant *Nitrospira*.

**Fig. 1.**
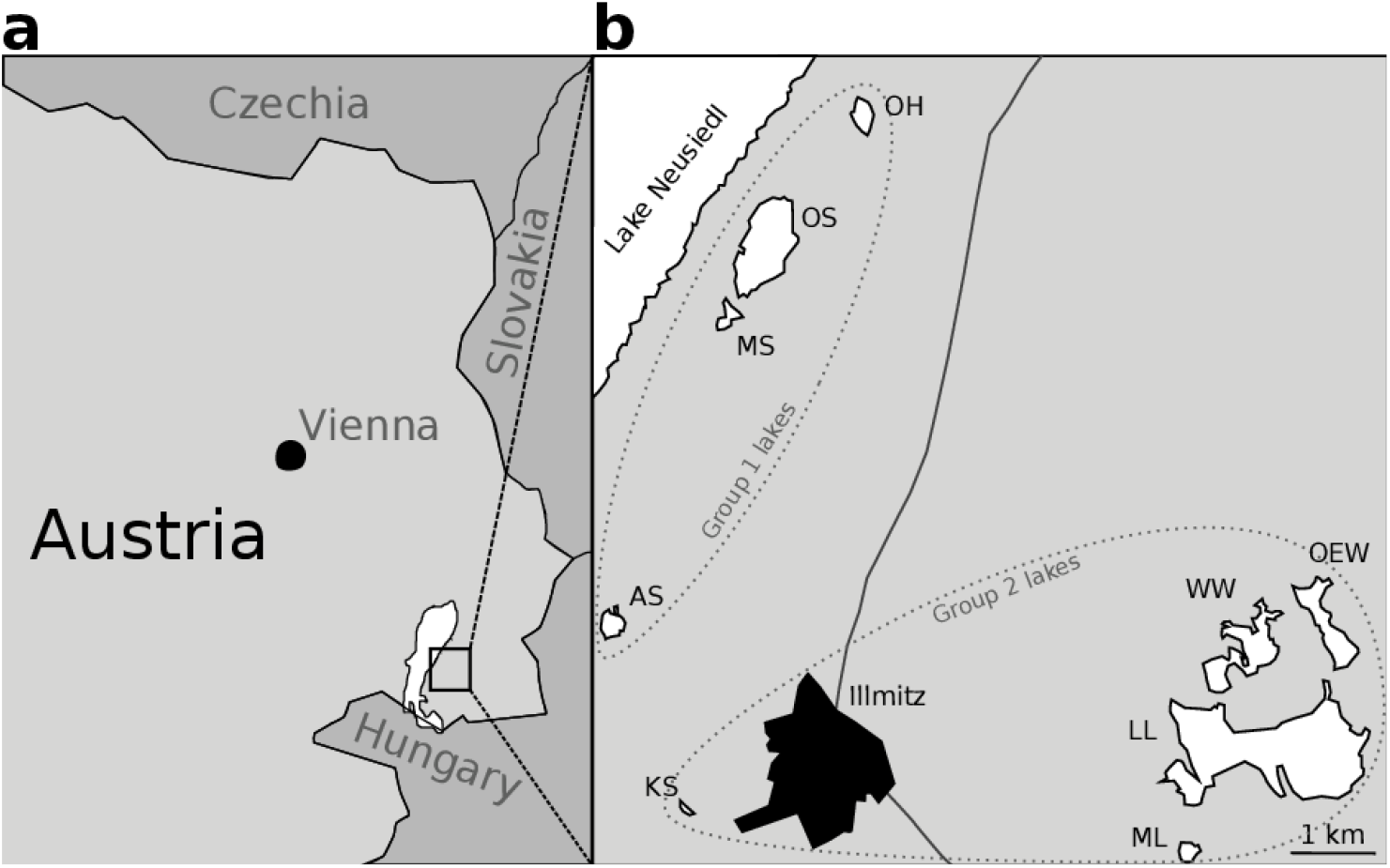
Map of the sampled aline-alkaline lakes. (a) Location of the sampling region in Austria. (b) Geographic location of the sampled lakes in the national park “Neusiedler See – Seewinkel”, Burgenland, Austria. The lakes are shown in white with the corresponding identifier abbreviations (see Table 1). Only lakes sampled for this study are shown. Dashed circles enclose lakes with similar *Nitrospira* communities (see Fig. 3).

## Materials and Methods

### *Sediment sampling and enrichment cultures of alkalitolerant* Nitrospira

Triplicate sediment samples (approx. 10 g) from nine saline-alkaline lakes in the national park “Neusiedler See - Seewinkel”, Burgenland, Austria (Fig. 1) were sampled from the top 10 cm of the sediments within a radius of five m in October 2013. The water pH and conductivity were measured for each lake at the time of sampling with a a SenTix 41 electrode and a WTW Multiline field instrument with a TetraCon 325, respectively. Salinity was inferred from conductivity based on a conversion factor, which had previously been established in experiments [31]. Dissolved organic carbon (DOC) concentrations were analyzed from sediment pore water filtered through precombusted glass fiber filters (GF/F, Whatman) and measured on a Total Carbon Analyzer (Sievers M9 Laboratory Analyzer, GE Analytical Instruments, Boulder, Colorado, U.S.A.). Nutrient concentrations (total nitrogen [TN], phosphorus-P, ammonium-N, nitrite-N, and nitrate-N) in lake waters were determined from sediment pore water using German Standard Methods [36]. Aliquots of both sediment and lake water samples were used as inoculum for nitrite oxidizer enrichment cultures, and the remaining material was stored at −20°C for molecular and chemical analyses. Concentration measurements of carbonate, total nitrogen, and trace metals in sediment samples were performed by the Austrian Agency for Health and Food safety (AGES GmbH, Vienna, Austria) according to their standard protocols. The results of the chemical measurements are listed in Table 1.

**Table 1.** Environmental properties determined for sediments, pore water (indicated by ^#^) and lake water (indicated by *) sampled in October 2013 from nine saline-alkaline lakes in the national park “Neusiedler See - Seewinkel”, Burgenland, Austria. Values marked with § denote data derived from the literature [32]. Missing data marked with NA (not available) was due to insufficient material for determination. Salinity values were estimated using a regionally constant correction factor [31]. Per cent units for carbonate and N_tot_ refer to the percentage weight of dried sediment. Abbreviations for lakes are as follows: AS, Albersee; MS, Mittlerer Stinker-See; OS, Oberer Stinker; OH, Obere Höll-Lacke; KS, Krautingsee; WW, Westliche Wörthenlacke; OEW, Östliche Wörthenlacke; LL, Lange Lacke; ML, Südliche Martinhoflacke.

| Lake (group) | AS (1) | MS (1) | OS (1) | OH (1) | KS (2) | WW (2) | OEW (2) | LL (2) | ML (2) |
| --- | --- | --- | --- | --- | --- | --- | --- | --- | --- |
| pH* | 9.98 | 9.77 | 10.3 | 9.82 | 9.71 | 9.18 | 9.16 | 9.42 | 8.9<sup>§</sup> |
| Conductivity [mS/ cm]* | 16.0<sup>§</sup> | 6.1<sup>§</sup> | 4.4<sup>§</sup> | 4.0<sup>§</sup> | 3.88 | 2.79 | 4.66 | 4.19 | 1.7<sup>§</sup> |
| Salinity [g/l]* | 12.8 | 4.88 | 3.52 | 3.2 | 3.1 | 2.23 | 3.73 | 3.35 | 1.36 |
| DOC [mg/l]<sup>#</sup> | NA | NA | NA | NA | 531.75 | 139.61 | 280.22 | 138.05 | NA |
| TN [mg/l]<sup>#</sup> | NA | NA | NA | NA | 33.54 | 4.09 | 15.75 | 5.69 | NA |
| NH<sub>4</sub><sup>+</sup>-N [µg/l]<sup>#</sup> | NA | 166 | NA | 1948 | 259 | 179 | 298 | 109 | 578 |
| NO<sub>2</sub><sup>-</sup>-N [µg/l]<sup>#</sup> | NA | 11 | NA | 398 | 30 | 4 | 13 | 7 | 12 |
| NO<sub>3</sub><sup>-</sup>-N [µg/l]<sup>#</sup> | NA | 179 | NA | 328 | 146 | 212 | 129 | 155 | 185 |
| PO<sub>4</sub><sup>-</sup>-P [µg/l]<sup>#</sup> | NA | 3100 | NA | 7850 | 133 | 58 | 1420 | 1840 | 774 |
| Carbonate [%] | 20.6 | 30.5 | 17.7 | 22.2 | 17.7 | 46.1 | 44.5 | 41.2 | 5.8 |
| N<sub>tot</sub> [%] | 0.087 | 0.109 | 0.092 | 0.081 | 0.135 | 0.293 | 0.585 | 0.164 | 0.032 |
| Fe [mol/gDW] | 7.48 | 10 | 10.16 | 10.33 | 14.97 | 12.06 | NA | 12.34 | 8.1 |
| Mn [mol/gDW] | 2.86 | 2.47 | 2.31 | 2.42 | 2.69 | 2.03 | NA | 3.3 | 7.25 |
| Cu [mol/gDW] | 0.1 | 0.21 | 0.08 | 0.12 | 0.1 | 0.15 | NA | 0.19 | 0.11 |
| Zn [mol/gDW] | 0.08 | 0.06 | 0.07 | 0.1 | 0.1 | 0.11 | NA | 0.29 | 0.05 |

Enrichment cultures of alkalitolerant NOB were established in mineral nitrite medium with a pH of 9 to 10.2 at 28°C. The medium was composed to reflect the chemical properties of the saline-alkaline lakes; however, trace elements were added as in Koch *et al*. [1]. The medium had the following composition: 1000 ml of distilled, millipore filtered water, 37 mg KCl, 53 mg CaCl_2_, 740 mg Na_2_SO_4_, 390 mg MgCl_2_, 150 mg KH_2_PO_4,_ 700 mg Na_2_CO_3_, 34 µg MnSO_4_ x H_2_O, 50 µg H_3_BO_3_, 70 µg ZnCl_2_, 72.6 µg Na_2_MoO_4_, x 2 H_2_O, 20 µg CuCl_2_ x 2 H_2_O, 24 µg NiCl_2_ x 6 H_2_O, 80 µg CoCl_2_ x 6 H_2_O, 1 mg FeSO_4_ x 7 H_2_O. The pH was monitored using indicator stripes (Macherey-Nagel) and a pH meter (WTW, Germany). Physiological tests were performed with selected *Nitrospira* enrichment cultures to determine their pH tolerance (with tested pH values ranging from 7.6 to 11) and nitrite concentration optimum of growth (with tested concentrations ranging from 0.15 to 1 mM NO_2_^−^). A detailed description of the cultivation procedure and physiological experiments is provided in the supplemental text.

### *Molecular analyses of* Nitrospira *community structures*

DNA extraction, PCR amplification, cloning, Illumina amplicon sequencing, and phylogenetic analyses of 16S rRNA gene and *nxrB* sequences, as well as rRNA-targeted fluorescence *in situ* hybridization (FISH), were performed as described in the supplemental text.

### *Metagenome sequencing*, Nitrospira *genome assembly, and analyses of genes putatively involved in haloalkalitolerance*

Cells of the “*Ca.* Nitrospira alkalitolerans” enrichment culture were harvested by centrifugation at 20.000 x g for 15 min and the cell pellet was used for DNA extraction according to Angel *et al.* [37]. Metagenome sequencing, assembly, binning, and annotation procedures are described in the supplemental text.

Specific genomic features of “*Ca.* N. alkalitolerans”, which are likely important for its adaptation to haloalkaline conditions, were identified by comparison to previously sequenced genomes of *Nitrospira* and *Nitrospina* that did not originate from haloalkaline habitats by using the OrthoFinder software [38] with default settings. Organisms used in these analyses were *Nitrospira moscoviensis [17]* and *N. defluvii* [39] (both canonical NOB), *N. inopinata* [18, 40] (moderately thermophilic comammox organism), “*Ca.* N. nitrosa” and “*Ca.* N. nitrificans” [19] (two mesophilic comammox organisms), and *Nitrospina gracilis* [41] (marine canonical nitrite oxidizer). Phylogenetic trees and the amino acid alignments of ATPase subunit *c* were reconstructed as described in the supplemental text.

## Data availability

The raw, demultiplexed amplicon sequencing datasets obtained in this study have been deposited at the European Nucleotide Archive (ENA) database under study accession number PRJEB34917. The raw metagenomic sequence reads obtained from the “*Ca.* N. alkalitolerans” enrichment culture have been deposited at ENA under study accession number PRJEB34830. The metagenome assembled (MAG) sequence and associated annotations of “*Ca.* N. alkalitolerans” are publicly available in MicroScope [42].

## Results and Discussion

### *Community composition of* Nitrospira *in the saline-alkaline lakes*

Members of the genus *Nitrospira* are the most diverse and widespread known NOB. However, reports of *Nitrospira* occurrence in alkaline habitats are scarce [23, 30], and a systematic assessment of their presence and activity in such extreme environments is missing. In this study, we discovered and investigated unusually alkalitolerant *Nitrospira* in saline-alkaline lakes of the national park “Neusiedler See – Seewinkel”, Burgenland, Austria using targeted amplicon profiling of the 16S rRNA gene and *nxrB*, of which the latter encodes the beta-subunit of nitrite oxidoreductase (the key enzyme for nitrite oxidation). In sediment samples from nine lakes, we detected phylogenetically diverse *Nitrospira* phylotypes which were affiliated with *Nitrospira* lineages I, II and IV (Fig. 2) [1].

**Fig. 2.**
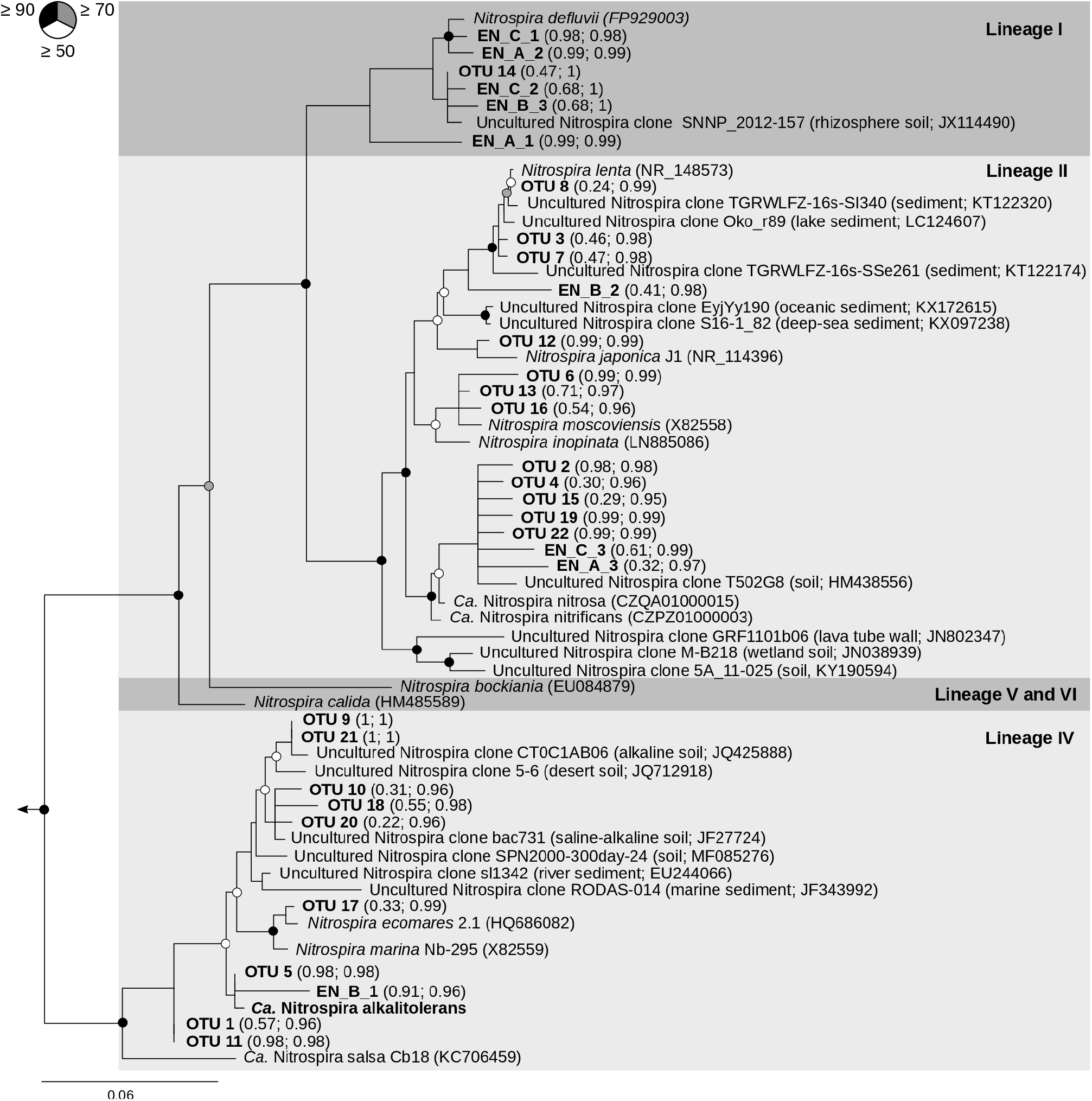
Phylogenetic maximum likelihood analysis based on the 16S rRNA gene sequences of selected representatives from the genus *Nitrospira* and of the *Nitrospira* members detected in sediments from nine saline-alkaline lakes. Sequences obtained in this study are printed in bold. “*Ca.* N. alkalitolerans” is the *Nitrospira* species cultured and further analyzed in this study. The tree was constructed using full length sequences and a 50% conservation filter resulting in 1310 valid alignment positions. Shorter sequences from this study, generated through amplicon and Sanger sequencing were added to the tree using the Evolutionary Placement Algorithm (EPA) without changing the overall tree topology. Numbers in brackets behind these sequences firstly denote the likelihood score of the exact placement and secondly the cumulative likelihood score of the placement within the cluster. Filled, grey, and open circles denote branches with ≥ 90%, ≥ 70% and ≥50% bootstrap support, respectively. *Leptospirillum ferrooxidans* (AJ237903), *Ca.* Magnetobacterium bavaricum (FP929063), *Thermodesulfovibrio yellowstonii* DSM 11347 (CP001147), and *Ca.* Methylomirabilis oxyfera (FP565575) were used as outgroup. The scale bar indicates 6% estimated sequence divergence.

The genomes of sequenced *Nitrospira* possess one to six paralogous copies of *nxrB*, and the *nxrB* copy numbers per genome remain unknown for the majority of uncultured *Nitrospira* [43]. This large variability likely affects relative abundance estimations of *Nitrospira* OTUs based on *nxrB* amplicon data. In contrast, all sequenced *Nitrospira* genomes contain only one ribosomal RNA (*rrn*) operon. Therefore, our further assessment of the *Nitrospira* community structures relies on the 16S rRNA gene amplicon datasets.

The estimated alpha-diversity of *Nitrospira* 16S rRNA gene phylotypes was compared across the nine examined lakes (Fig. S2). The inverse Simpson’s index of the *Nitrospira* communities was negatively correlated with pH and the nitrite concentration (p = 0.00004, Tau-b = −0.53 for pH and p = 0.03, Tau-b = −0.36 for nitrite). The decrease of *Nitrospira* diversity with increasing pH may indicate that only specific *Nitrospira* phylotypes tolerate highly alkaline conditions.

The *Nitrospira* communities clustered into two distinct major groups (Fig. 3). Group 1 mainly comprised the communities from those lakes, which are located closely to the shore of the much larger Lake Neusiedl, whereas group 2 contained the communities from the remaining lakes that are farther away from Lake Neusiedl (Fig. 1). The average pH and salinity in the water of lakes from the group 1 cluster were 9.97 ± 0.24. and 6.1 ± 4.1 g/l, respectively. These values were significantly higher (Welch’s t-test; p = 0.00001 for pH and p = 0.017 for salinity) than the mean pH of 9.37 ± 0.26 and salinity of 2.74 ± 0.88 g/l in the group 2 lakes (Table 1). None of the other determined lake properties at time of sampling differed significantly between the two groups. The *Nitrospira* phylotypes with the highest relative abundance in the sediments from group 1 were OTU1 and OTU20, both affiliated with *Nitrospira* lineage IV, whereas these OTUs were nearly absent from the sediments of the lakes in group 2 (Fig. 3). In contrast, the predominant phylotypes in the group 2 lake sediments were affiliated with *Nitrospira* lineage II (Fig. 3). Consistent with these results, a principal coordinate analysis showed a clear separation of the *Nitrospira* communities with the same two groups separated on the first axis of the ordination (Fig. S3). These results indicate a strong influence of pH and salinity on the composition of the *Nitrospira* communities. Members of *Nitrospira* lineage IV are adapted to saline conditions and are commonly found in marine ecosystems [15, 44–48]. However, to date no *Nitrospira* species have been described to tolerate elevated pH conditions. Our results show that a substantial diversity of *Nitrospira* is able to colonize alkaline environments. The data also indicate a niche differentiation between lineages IV and II in saline-alkaline lakes, which likely includes a higher tolerance of the detected lineage IV organisms towards an elevated pH and salinity.

**Fig. 3.**
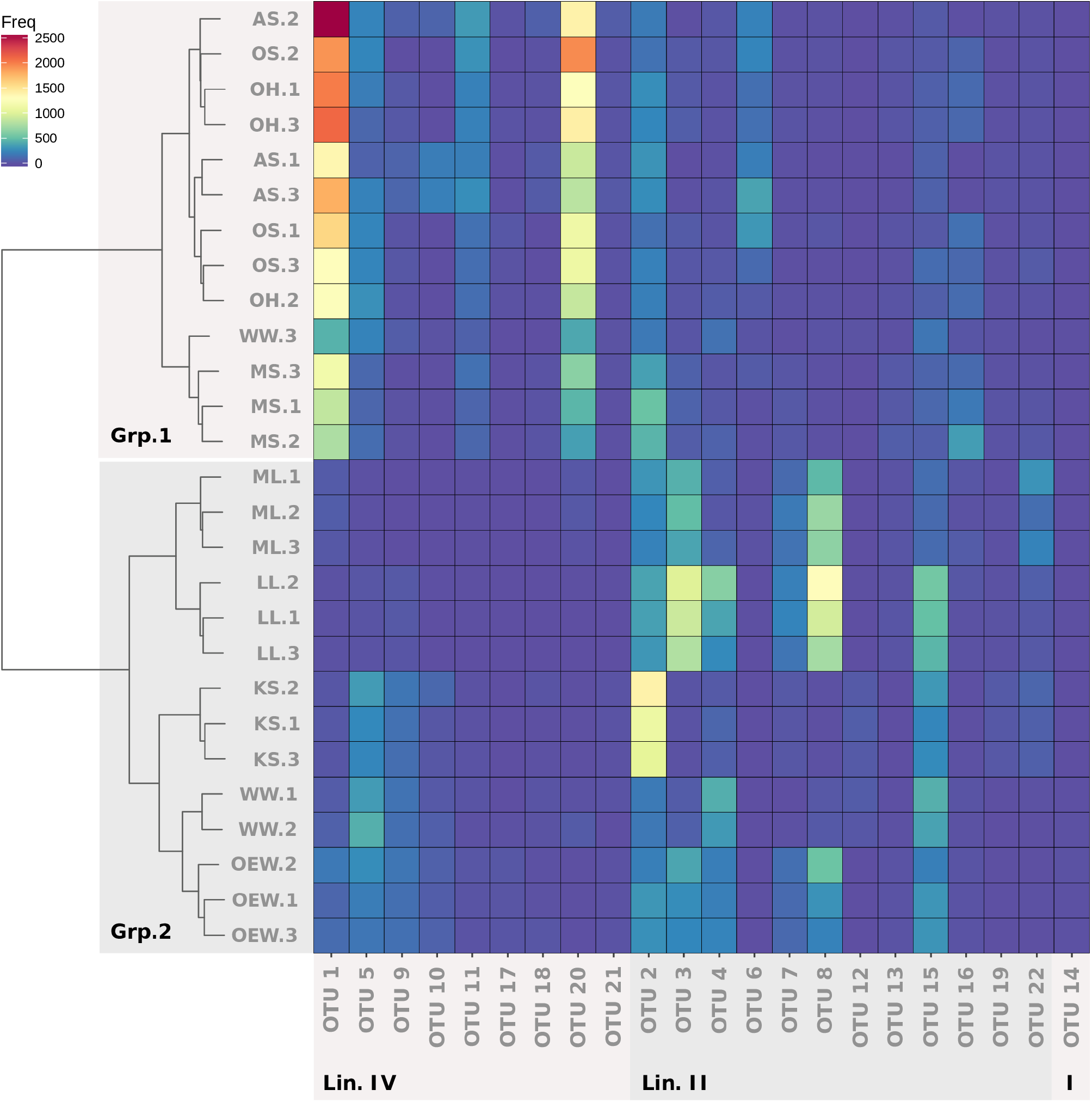
Normalized abundances of *Nitrospira* 16S rRNA gene phylotypes detected in triplicate sediment samples from nine saline-alkaline lakes. *Nitrospira* communities are grouped by hierarchical clustering on the *y*-axis, and OTUs are grouped by phylogenetic affiliation on the *x*-axis. Lake names are abbreviated as in Table 1. Lin. IV, *Nitrospira* lineage IV; Lin. II, *Nitrospira* lineage II; I, *Nitrospira* lineage I; Freq, normalized frequency counts; Grp.1, group 1 lakes; Grp.2, group 2 lakes (see also Fig. 1).

### *Metagenome sequencing and physiology of alkalitolerant* Nitrospira *enrichments*

Following the inoculation of mineral nitrite medium flasks with sediment and/or water samples from four saline-alkaline lakes (LL, WW, KS and OEW; abbreviations see Table 1), we initially obtained 17 enrichment cultures that oxidized nitrite to nitrate. Based on FISH analyses with *Nitrospira*-specific 16S rRNA gene-targeted probes and Sanger sequencing of cloned 16S rRNA genes, several of these preliminary enrichment cultures contained co-existing phylotypes from *Nitrospira* lineages I, II, and IV as well as from the genus *Nitrobacter* (data not shown). Members of the genera *Nitrotoga* and *Nitrospina* were screened for by FISH or PCR, but were not detected.

We used three of the enrichments which contained only *Nitrospira* NOB and originated from different lakes (referred to as EN_A from lake OEW, EN_B from lake LL, and EN_C from lake WW comprising approx. 35% *Nitrospira* in relation to the total microbial community based on FISH analysis) to determine the pH range for activity of the enriched *Nitrospira* members. Enrichment cultures EN_A and EN_C contained phylotypes from *Nitrospira* lineages I and II, while EN_B contained phylotypes from lineages I, II, and IV as determined by 16 rRNA gene amplicon cloning and Sanger sequencing (Fig. 2). The continued presence of these *Nitrospira* phylotypes for more than two years, despite several serial dilution transfers, demonstrates their tolerance to the alkaline incubation conditions and suggests that they were native to the saline-alkaline environment which they were sampled from. Hence, we conclude that at least the highly similar uncultured *Nitrospira* OTUs detected by amplicon sequencing (Fig. 2) were most likely also native inhabitants of the saline-alkaline lakes. Aliquots of each enrichment culture were incubated with nitrite as the sole added energy source for six weeks at pH 7.61-7.86 and 9-9.04, respectively. During this period, pH had no significant effect on nitrite utilization (Pearson correlation coefficient ≥ 0.96 with, p ≤ 0.01 for all three enrichments) and nitrate production (Pearson correlation coefficient ≥ 0.98 with, p ≤ 0.01 for all three enrichments) over time for any of the three enrichments (Fig. S4). Subsequently, the enrichment culture aliquots that had been incubated at pH 9-9.04 were sequentially incubated at pH 9.97-10, 10.24-10.52, and 10.72-11.02 for eight to nine days at each pH (Table S3). For all three enrichments, the observed nitrate production tended to be slower at pH 9.97-10 and 10.24-10.52 than at pH 9-9.04 (Fig. S4 and S5). At pH 10.72-11.02, no nitrite consumption was detected (Fig. S5). The trends observed at pH 10.24-10.52 and above were in stark contrast to the persistently high nitrite-oxidizing activity of the enrichments when routinely cultured at pH 9-10 for several weeks. While it was not possible to determine based on our data whether all *Nitrospira* phylotypes present in the three enrichments responded equally to the tested pH conditions, we can conclude that the activity of at least some *Nitrospira* remained unaffected up to pH 9 and had an upper limit between pH 10.5 and 10.7. This is remarkable, because previously enriched or isolated *Nitrospira* strains were not cultivated above pH 8.0 except for two *Nitrospira* cultures from geothermal springs, which showed activity up to pH 8.8 [4] or pH 9.0 [7]. To our knowledge, this is the first report of nitrite oxidation by *Nitrospira* at pH values above 9 and as high as 10.5.

Further analyses focused on one additional enrichment, which had been inoculated with sediment from lake Krautingsee, belonging to the group 2 of the analyzed lakes (KS, Table 1). In contrast to the other enrichment cultures, this enrichment contained only lineage IV *Nitrospira* based on FISH analysis (Fig. 4a). *Nitrospira*-specific, 16S rRNA gene and *nxrB*-targeted PCR and phylogeny detected one phylotype from *Nitrospira* lineage IV that was related to other phylotypes detected from the lakes, specifically OTU 5 and EN_B_1 (16S rRNA gene, 100% and 98% nucleotide sequence identity, respectively; Fig. 2) and OTU 2 (*nxrB*, 98.5% nucleotide sequence identity; Fig. S1). The OTU 5 phylotype occurred in most of the analyzed lakes (Fig. 3). Thus, the closely related enrichment from lake KS may represent *Nitrospira* that could adapt to a relatively broad range of conditions, while some of the other OTUs were more abundant in specific lakes only (Fig. 3). The enriched *Nitrospira* reached a high relative abundance in the enrichment culture of ~60% of all bacteria based on metagenomic read abundance (see below) and observation by FISH.

**Fig. 4.**
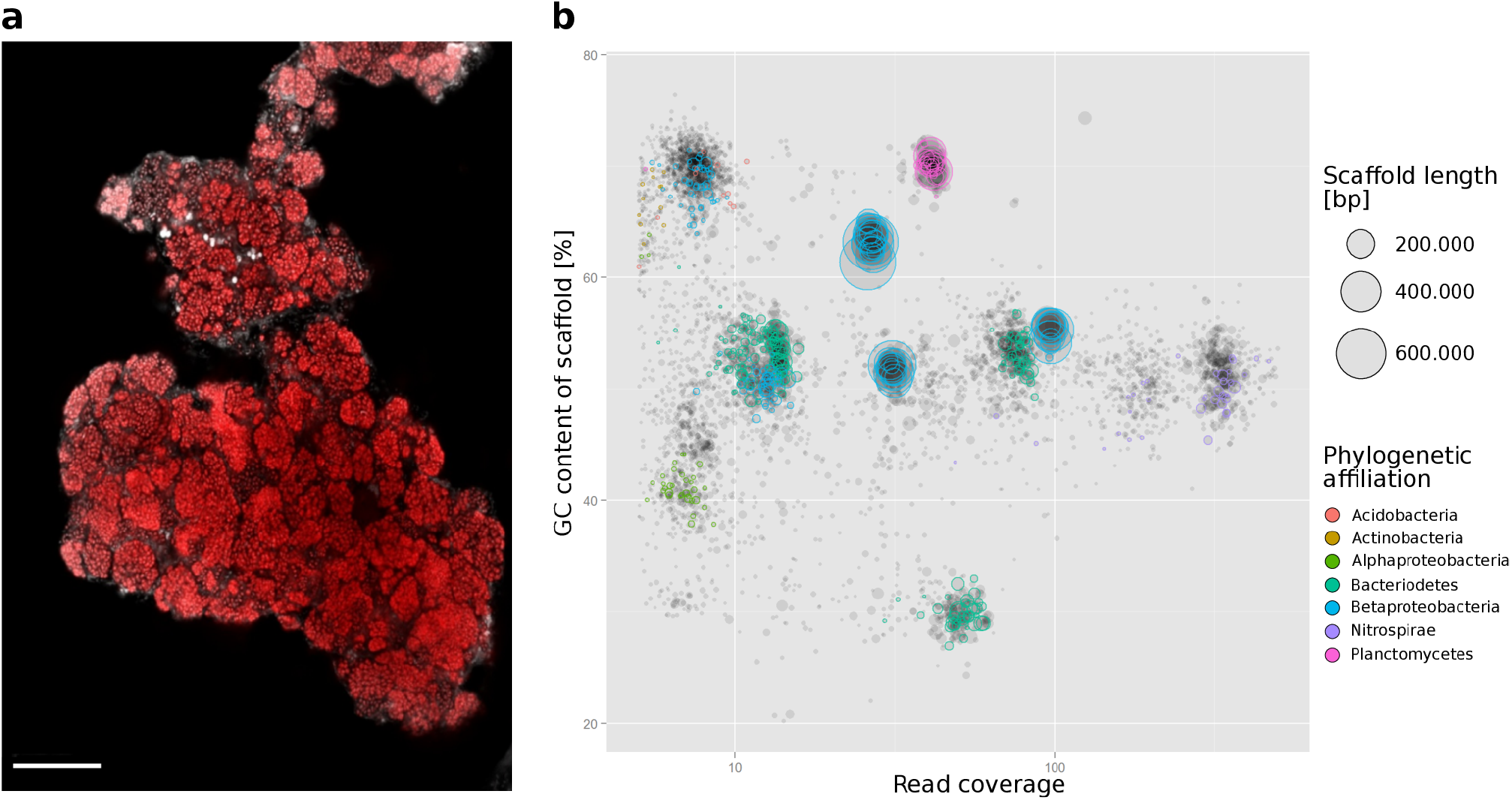
Visualization and metagenomic analysis of the “*Ca.* N. alkalitolerans” enrichment. (a) FISH image showing dense cell clusters of “*Ca.* N. alkalitolerans” in the enrichment culture. The “*Ca.* N. alkalitolerans” cells appear in red (labelled by probe Ntspa1151 which has 1 mismatch at the 3’ end to the 16S rRNA gene sequence of “*Ca.* N. alkalitolerans”; the absence of lineage II *Nitrospira* in the enrichment culture was confirmed by the application of the competitor oligonucleotides c1Ntspa1151 and c2Ntspa1151 as indicated in the supplemental text). Other organisms were stained by DAPI and are shown in light grey. Scale bar, 25 µm. (b) Phylogenetic affiliation of the metagenome scaffolds from the “*Ca.* N. alkalitolerans” enrichment, clustered based on sequence coverage and the GC content of DNA. Closed circles represent scaffolds, scaled by the square root of their length. Clusters of similarly colored circles represent potential genome bins.

High-throughput metagenome sequencing, scaffold assembly, and binning revealed that the enrichment contained three *Nitrospira* strains that could be separated into three genome bins based on sequence coverage data (Table S1, Fig. S7). No other NOB were identified in the metagenome, and the three *Nitrospira* bins represented the most abundant organisms in the enrichment culture (Fig. 4b). Since the genome-wide average nucleotide identity (gANI) values were above the current species threshold of 95% [49] (Table S1), the three bins likely represented very closely related strains of the same *Nitrospira* lineage IV species with unique genetic components. From the predominant (based on coverage data) *Nitrospira* sequence bin, an almost complete metagenome-assembled genome (MAG) was reconstructed, which met the criteria for a “high-quality draft” genome [50] (Table S1), and used for comparative genomic analysis. Genome-wide, pairwise comparison of the gANI and average amino acid (gAAI) identity between this MAG and *Nitrospira marina* as the only other genome-sequenced and cultured *Nitrospira* lineage IV representative resulted in values of 80.1 and 77.3, respectively. The 16S rRNA gene, which had been retrieved from the MAG, was 97.90% identical to the 16S rRNA gene of *N. marina*, 97.87% identical to “N. strain Ecomares 2.1”, 94.92% to “*Ca*. N. salsa”, and 94.51% to “*Nitrospira* strain Aa01”, which are the other cultured members of *Nitrospira* lineage IV [15, 44, 47, 48]. These values are below the current species threshold of 98.7-99% for 16S rRNA genes [54]. Based on the low gANI and 16S rRNA gene sequence identities to described *Nitrospira* species, and additionally considering the distinct haloalkalitolerant phenotype (see also below), we conclude that the enriched *Nitrospira* represent a new species and propose “*Ca*. Nitrospira alkalitolerans” as the tentative name.

The enrichment culture was maintained at a pH of 9 to 10 and a salt concentration of 2 g/l, resembling the natural conditions in the saline-alkaline lakes based on available data from five years. “*Ca*. N. alkalitolerans” grew in dense flocks (Fig. 4a), thereby possibly relieving the pH stress [51]. Its nitrite-oxidizing activity was not affected when the pH in the cultivation medium decreased below 8. However, no nitrite oxidation was observed when the enrichment culture was transferred into medium with 4× to 8× higher salt concentrations, the latter resembling marine conditions. Thus, “*Ca.* N. alkalitolerans” is best described as a facultatively haloalkalitolerant organism that oxidizes nitrite as an energy source over a wide range of pH and under hyposaline conditions. This phenotype is certainly advantageous in the investigated saline-alkaline lakes, as these lakes are prone to evaporation in summer, which causes a temporarily elevated salinity and alkalinity in the remaining water body and the sediment [35].

The enrichment culture of “*Ca.* N. alkalitolerans” oxidized nitrite over a broad range of initial nitrite concentrations tested, although an extended lag phase of 10 to 15 days occurred at the higher concentrations of 0.7 and 1 mM nitrite (Fig. S6). Similarly, a lag phase at elevated nitrite concentrations was also observed for the *Nitrospira* lineage II member *Nitrospira lenta* [52]. A preference for low nitrite levels is consistent with the presumed ecological role of nitrite-oxidizing *Nitrospira* as slow-growing K-strategists, which are adapted to low nitrite concentrations [52–54].

### Genomic adaptations to the saline-alkaline environment

As described below, comparative genomic analysis of “*Ca.* N. alkalitolerans” revealed several features that distinguish this organism from other known NOB and likely form the basis of its tolerance towards elevated alkalinity and salinity (Fig. 5).

**Fig. 5.**
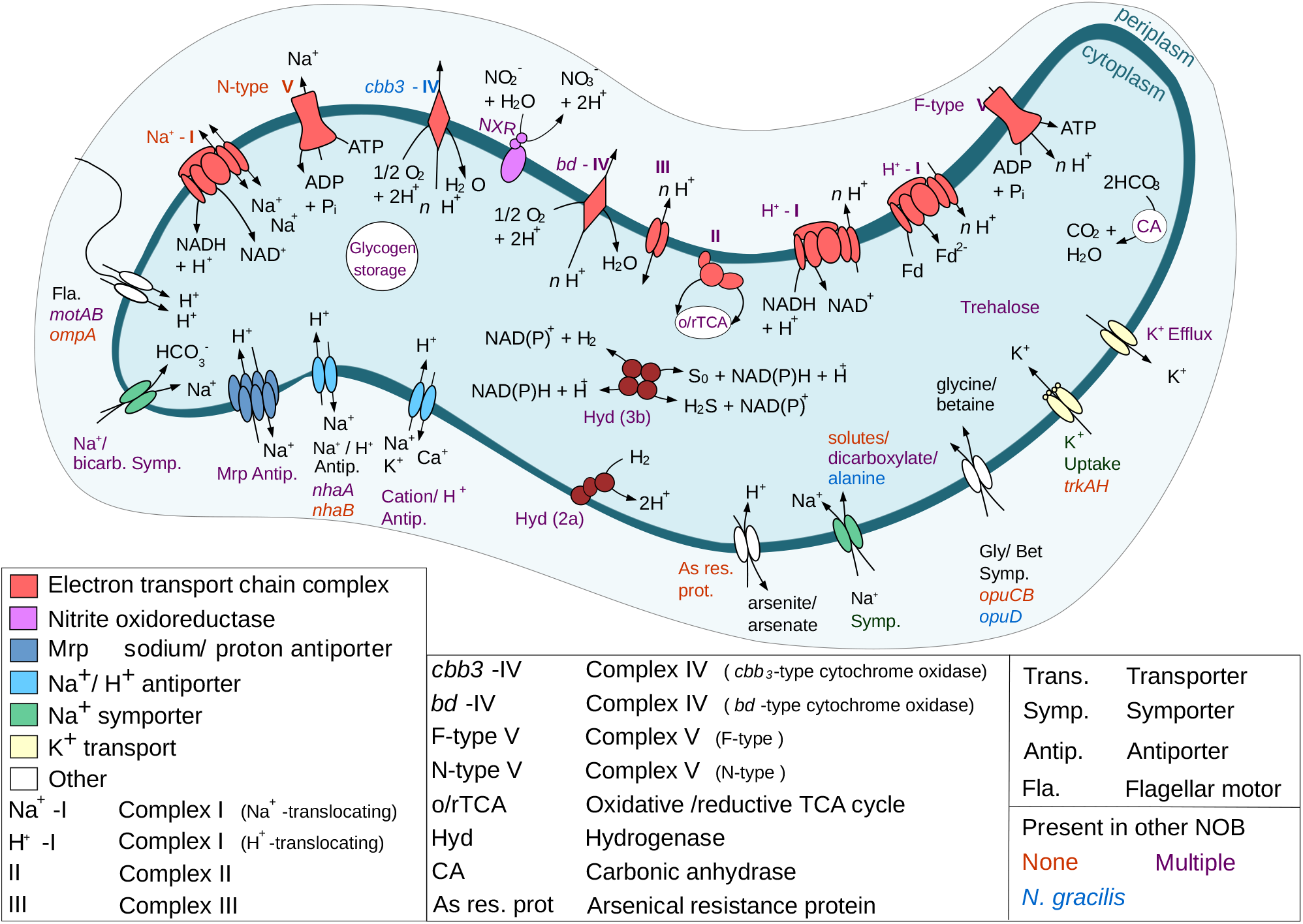
Cell metabolic cartoon constructed from the genome annotation of “*Ca.* N. alkalitolerans”. Features putatively involved in the adaptation to a high alkalinity and salinity, and selected core metabolic pathways of chemolithoautotrophic nitrite-oxidizing *Nitrospira*, are shown. Note that the transport stoichiometry of the ion transporters in “*Ca.* N. alkalitolerans” remains unknown. Colors of text labels indicate whether adaptive features are present (*i.e.*, have homologs) in the genomes of other NOB (red, feature is not present in any other characterized NOB; blue, feature is present only in the marine *Nitrospina gracilis*; purple, feature is present in several other characterized NOB).

#### (i) Cytoplasmic pH and ion homeostasis

At high pH, alkaliphilic and alkalitolerant microbes maintain a higher transmembrane electrical potential (ΔΨ) component of the proton motive force (PMF) than usually found in neutrophiles. The high ΔΨ is required to maintain PMF, because the ΔpH component of the PMF is reversed when the extracellular pH is higher than the intracellular pH [55]. Like in neutrophiles, the ΔΨ of alkaliphiles is negative inside the cell relative to the outside [55]. Furthermore, the intracellular pH must be kept below the (extremely) alkaline extracellular pH. At elevated salinity, resistance against high salt concentrations is an additional, fundamental necessity for survival. All this requires a tightly regulated pH and ion homeostasis, in which cation transmembrane transporters play key roles [55–57]. The “*Ca.* N. alkalitolerans” genome codes for various Na^+^-dependent transporters (Fig. 5, Table S2) including secondary Na^+^/H^+^ antiporters that are involved in pH homeostasis in other organisms: two copies of a group 3 Mrp-type Na^+^/H^+^ antiporter [58, 59] encoded by the seven genes *mrpA-G*, and monovalent cation-proton antiporters of the types NhaA and NhaB, each of which is encoded by a single gene [60]. The Mrp antiporter is crucial for growth at high pH and elevated salinity in alkaliphilic *Halomonas* spp. and *Bacillus* spp., where it exports Na^+^ and imports H^+^, thus contributing to the maintenance of a lower intracellular pH compared to the environment (*e.g.*, cytoplasmic pH 8.3 at external pH ~10.5) [50 and references cited therein, 56]. The Mrp proteins may form a large surface at the outside of the cytoplasmic membrane that could support proton capture under alkaline conditions [55, 58]. Nha-type antiporters are widely distributed among non-extremophilic and extremophilic organisms [56]. Being involved in the homeostasis of Na^+^ and H^+^, they are important for survival under saline and/or alkaline conditions [57]. In *E. coli*, NhaA is regulated by the cytoplasmic pH and it catalyzes the import of 2H^+^ with the concurrent export of one Na^+^. This electrogenic activity is driven by ΔΨ and maintains pH homeostasis at elevated external pH [52 and references cited therein]. The simultaneous presence of the two antiporters NhaA and NhaB has been associated with halophilic or haloalkaliphilic phenotypes in other organisms [56, 60]. Although the regulation and cation transport stoichiometry of the homologs in “*Ca.* N. alkalitolerans” remain unknown, the Mrp- and Nha-family antiporters most likely exhibit important physiological roles in this organism and support its survival under haloalkaline conditions. Possibly, “*Ca.* N. alkalitolerans” can even combine its growth in dense flocks with the extrusion of protons by its numerous proton transporters thereby lowering the pH inside the flock [51].

One of the two *nhaB* genes present in the “*Ca.* N. alkalitolerans” genome is located in an interesting genomic region that also contains all genes encoding the group 3 Mrp-type Na^+^/ H^+^ antiporter (Fig. S8). The two genes downstream from *mrpD* display sequence similarity to the NADH dehydrogenase (complex I) subunits NuoM and NuoL. However, based on the genomic context they are more likely additional *mrpA*- and/or *mrpD*-like genes, as these Na^+^/ H^+^ antiporter subunits are evolutionary related to NuoM and NuoL [62]. Multiple copies of subunits NuoM and NuoL of the NADH dehydrogenase are encoded elsewhere in the genome, partially in larger *nuo* operons (see Table S2). Moreover, the locus contains one gene coding for the low-affinity, high flux Na^+^/HCO_3_^−^ uptake symporter BicA [63] and gene *motB* encoding a H^+^-translocating flagellar motor component (Fig. S8). In the haloalkalitolerant cyanobacterium *Aphanothece halophytica*, a similar clustering of *bicA* with genes coding for Na^+^/H^+^ antiporters has been described. The authors proposed a model of cooperation between these transporters, where Na^+^ extruded by the Na^+^/H^+^ antiporters could drive the uptake of HCO_3_^−^ by BicA under alkaline conditions when CO_2_ becomes limiting [64]. Sodium-driven import of HCO^3^ could be an essential feature for “*Ca.* N. alkalitolerans”, because bicarbonate is the main source of inorganic carbon for autotrophic organisms, but becomes less accessible at high pH >10 [56]. A carbonic anhydrase, which is also present in the genome (Fig. 5, Table S2), can convert the imported HCO_3_^−^ to CO_2_ for carbon fixation *via* the reductive tricarboxylic acid cycle (Fig. 5).

Since cytoplasmic K^+^ accumulation may compensate for Na^+^ toxicity at elevated intracellular pH [65], many alkaliphiles retain an inward directed K^+^ gradient [56]. The potassium uptake transporters of the Trk family contribute to pH and K^+^ homeostasis of halo- and/or alkaliphiles [56]. TrkAH catalyzes the NAD^+^-regulated uptake of K^+^ possibly coupled with H^+^ import [66]. Moreover, kinetic experiments revealed that TrkAH of the gammaproteobacterium *Alkalimonas amylolytica* is salt-tolerant and functions optimally at pH >8.5 [67]. “*Ca.* N. alkalitolerans” encodes a TrkAH complex (Fig. 5, Table S2), which may be a specific adaptation to its haloalkaline environment as no homologous K^+^ transporter has been identified yet in any other NOB genome. Under more neutral pH conditions, Kef-type K^+^ efflux pumps, which are present in two copies in the “*Ca.* N. alkalitolerans” genome, could excrete excess K^+^ (Fig. 5, Table S2).

#### (ii) Adaptations of the energy metabolism

Aside from the different cation transporters (see above), “*Ca.* N. alkalitolerans” also encodes several mechanisms for cation homeostasis that are linked to membrane-bound electron transport and energy conservation. Like in other aerobic alkaliphiles [57], ATP synthesis is likely catalyzed by a canonical, H^+^-translocating F_1_F_O_-ATPase (Fig. 5, Table S2). In addition, the genome contains all genes of a predicted Na^+^-translocating N-ATPase [68] (Fig. 5, Fig. S9, Table S2). N-ATPases form a separate subfamily of F-type ATPases and have been suggested to be ATP-driven ion pumps that extrude Na^+^ cations [68] or H^+^ [69]. The *c* subunit of the N-ATPase in the genome of “*Ca.* N. alkalitolerans” contains the typical amino acid motifs for Na^+^ binding and transport [68] (Fig. S10). Subunits *a* and *c* of the N-ATPase, which are involved in ion transport, are most similar to homologs from the halotolerant, sulfate-reducing *Desulfomicrobium baculatum* (81.5% AA identity) and the haloalkalitolerant, sulfur-oxidizing *Sulfuricella denitrificans* (88.2% AA identity), respectively. Hence, in “*Ca.* N. alkalitolerans”, the N-ATPase may contribute to the maintenance of ΔΨ, the generation of a sodium motive force (SMF), and salt resistance (Fig. 5).

The genome of “*Ca.* N. alkalitolerans” encodes two different types of NADH:quinone oxidoreductase (complex I of the electron transport chain) (Fig. 5, Table S2). Firstly, the organism possesses all 14 genes of type I NADH dehydrogenase (*nuoA* to *nuoN*). They are present in one to three copies each. The *nuo* genes are mostly clustered at several genomic loci (Table S2) and are most similar to either of the two *nuo* operons present in *Nitrospira defluvii* [39], with AA identities between 41% and 90%. As mentioned above, *nuoL/M*-like genes at loci without other *nuo* genes might represent subunits of cation antiporters.

The genome furthermore contains a locus encoding all six subunits of a Na^+^-dependent NADH:quinone oxidoreductase (Nqr or type III NAD dehydrogenase) (Fig. 5, Table S2). The locus is situated on a single contig in the vicinity of transposase genes, indicating that “*Ca*. N. alkalitolerans” might have received this type of complex I by lateral gene transfer. The gene of subunit E, which takes part in Na^+^ translocation [70], is most similar to a homolog in the ammonia-oxidizing bacterium *Nitrosomonas nitrosa* (86% AA identity).

The metabolic model for *N. defluvii* [39] assumes that two different versions of the H^+^-dependent complex I (Nuo) are used for forward or reverse electron transport, respectively. *Nitrospira* possess a canonical Nuo that is likely used for PMF generation during the forward flow of low-potential electrons from the degradation of intracellular glycogen or from hydrogen as an alternative substrate (see also below). In addition, reverse electron transport is essential in NOB to generate reducing power for CO_2_ fixation. In *Nitrospira*, a second (modified) form of Nuo with duplicated proton-translocating NuoM subunits might use PMF to lift electrons from quinol to ferredoxin [71]. The reduced ferredoxin is required for CO_2_ fixation *via* the rTCA cycle. As expected, “*Ca.* N. alkalitolerans” possesses these two Nuo forms that are conserved in other characterized *Nitrospira* members. In addition, the Na^+^-dependent Nqr complex might function in two directions in “*Ca*. N. alkalitolerans” as well. During forward electron flow, Nqr would contribute to SMF generation (Fig. 5). Reverse operation of the Nqr could generate NADH while importing Na^+^, thus utilizing SMF for the reduction of NAD^+^ with electrons derived from quinol (Fig. 5). Hence, the two types of complex I are likely involved in essential electron transport and the fine-tuning of PMF and SMF. They probably cooperate with the Na^+^- and the H^+^-translocating ATPases and the various cation transporters (see above) to adjust the cytoplasmic ion concentrations and the membrane potential in response to the environmental salinity and pH.

In addition to a novel “*bd*-like” cytochrome c oxidase, which is commonly found in *Nitrospira* genomes [16, 39], the genome of “*Ca.* N. alkalitolerans” contains a locus with fused genes for a *cbb_3_*-type cytochrome c oxidase (Fig. 5, Table S2) similar to the one present in the marine nitrite oxidizer *Nitrospina gracilis* [41]. The c*bb_3_*-type terminal oxidases usually exhibit high affinities for O_2_ [72] and may allow “*Ca*. N. alkalitolerans” to sustain respiration at low oxygen levels.

Interestingly, “*Ca*. N. alkalitolerans” encodes two different hydrogenases and the accessory proteins for hydrogenase maturation (Fig. 5, Table S2). First, it possesses a group 2a uptake hydrogenase that is also found in *N. moscoviensis*, which can grow autotrophically on H_2_ as the sole energy source [16]. Second, “*Ca*. N. alkalitolerans” codes for a putative bidirectional group 3b (sulf)hydrogenase that also occurs in other NOB and in comammox *Nitrospira* [18, 41] but has not been functionally characterized in these organisms. Experimental confirmation of H_2_ utilization as an alternative energy source and electron donor by “*Ca*. N. alkalitolerans” is pending. However, we assume that this capability would confer ecophysiological flexibility, especially if nitrite concentrations fluctuate and H_2_ is available at oxic-anoxic boundaries in biofilms or upper sediment layers. While electrons from the group 2a hydrogenase are probably transferred to quinone [16], the group 3b hydrogenase might reduce NAD^+^ [41] and fuel forward electron transport through the Nuo and Nqr complexes (see above).

#### (iii) Osmoadaptation

The intracellular accumulation of compatible solutes is an important mechanism allowing microorganisms to withstand the high osmotic pressure in saline habitats [56]. “*Ca.* N. alkalitolerans” has the genetic capacity to synthesize or import the compatible solutes trehalose, glycine betaine, and glutamate (Fig. 5). For trehalose synthesis the gene *treS* of trehalose synthase (Table S2), which enables trehalose synthesis from maltose, is present. The genes *opuD* and *opuCB* for glycine betaine import (Table S2) have been identified in the marine *Nitrospina gracilis* [41], but not yet in any *Nitrospira* species. For glutamate synthesis, the genes *gltB* and *gltD* were identified (Table S2). They code for the alpha and beta subunits of glutamate synthase, which catalyzes L-glutamate synthesis from L-glutamine and 2-oxoglutarate with NADPH as cofactor. In addition, we identified adaptations of “*Ca.* N. alkalitolerans” to the low availability of iron and the presence of toxic arsenite in saline-alkaline systems (supplemental text).

## Conclusions

This study shows that diverse *Nitrospira* phylotypes are able to colonize saline-alkaline lakes, and that members of these lineages can carry out chemolithoautotrophic nitrite oxidation under strongly alkaline conditions up to pH 10.5. The genomic analysis of the newly cultured *Nitrospira* species “*Ca.* Nitrospira alkalitolerans” has revealed several adaptive features, many of which are also found in other haloalkalitolerant or –philic microorganisms but are missing in other characterized NOB. These results extend our picture of nitrogen cycling in extreme habitats and push the known limits of nitrite oxidation to an unusually high pH. The presence of hydrogenase genes in “*Ca.* N. alkalitolerans” suggests that alkalitolerant NOB can utilize alternative energy metabolisms and thus share at least part of the physiological versatility known from their neutrophilic relatives [13, 16, 17, 73]. As a next step it will be crucial to determine which ammonia oxidizers and/or comammox organisms coexist with alkalitolerant NOB and drive nitrification in saline-alkaline ecosystems.

## Acknowledgements

The authors thank Beate Pitzl from the Biological Station WasserCluster Lunz, Austria for chemical analysis and Zsófia Horváth for help with sample collection. We are further grateful to Sebastian Lücker for developing and sharing a *Nitrospina nxrB* reverse primer sequence. Queralt Güell-Bujon is acknowledged for maintaining and optimizing growth conditions of enrichments. This research was supported by the Austrian Science Fund (FWF) grants T938 (to AD), and P25231-B21 and P27319-B21 (both to HD).

## Author contributions

AD and HD conceived the study. AD took the samples, performed the physiological experiments and analyzed the data. AD, KK, HK, MS, JS set up the enrichments and collected data. CWH, SMK and MA processed the amplicon and metagenomic raw data. AD performed phylogenetic and comparative genomic analyses. AD and HD wrote the manuscript with input from all authors.

## Supplemental Text

### Supplemental Materials and Methods

#### Enrichment of alkalitolerant Nitrospira species

Nitrite-oxidizing enrichment cultures were established by inoculation of 40 ml sterile, mineral nitrite medium containing 0.5 mM filter-sterilized NaNO_2_ with approx. 0.1 g of fresh sediment or 1 ml of lake water. Enrichment cultures were incubated without agitation in 100 ml glass bottles in the dark at 28°C. Freshly prepared medium had a pH of 10.2 that decreased to 9 during approximately two months of cultivation. Actively nitrite-oxidizing enrichment cultures were further purified and propagated by 1:100 dilution in fresh nitrite medium. The nitrite and nitrate concentration in the medium were regularly checked by using nitrite/nitrate test stripes (Macherey-Nagel), and nitrite was repeatedly replenished when completely consumed. Precise chemical measurements [2] of the nitrite and nitrate concentrations were regularly performed during the early stages of the *Nitrospira* enrichment to confirm that nitrite was stoichiometrically oxidized to nitrate.

#### Fluorescence in situ hybridization (FISH) and microscopy

The abundance of *Nitrospira* in the enrichment cultures was regularly monitored by FISH with 16S rRNA gene-targeted probes that were doubly labeled with the fluorochromes Cy3, Cy5, or FLUOS. Probes targeting most bacteria (EUB338 probe mix at a formamide (FA) concentration of 35%; [3, 4], the phylum Nitrospirae (probe Ntspa712 with competitor at a FA concentration of 35%; [5]) and most lineage II plus some lineage IV species of the genus Nitrospira (probe Ntspa1151 at a FA concentration of 35%; [6]), the genus *Nitrobacter* (probe Nit3 with competitor at a FA concentration of 35%; [7]), the genus *Nitrotoga* (probe Ntoga122 with two competitors at a FA concentration of 40%; [8]) and probe NON338 [9] as a control for nonspecific probe binding were applied. To specifically detect lineage II *Nitrospira* only, probe Ntspa1151 was used in equimolar concentrations at a FA concentration of 35% with two newly designed, unlabeled competitor oligonucleotides targeting lineage IV *Nitrospira* (c1Ntspa1151: 5‘-TTA TCC TGG GCA GTC TCT CC-3’ and c2Ntspa 1151: 5‘-TTA TCC TGG GCA GTC TCT TC-3’). FISH was combined with nonspecific fluorescent labeling of all cells by 4’,6’-diamidino-2-phenylindole (DAPI). For FISH, aliquots of the enrichment cultures were formaldehyde-fixed and FISH was performed according to standard protocols [10]. Fluorescence micrographs of probe-stained organisms were acquired with an inverted Leica TCS SP8X confocal laser scanning microscope that was equipped with a 405 nm UV diode, a Leica supercontinuum white light laser, two photomultiplier (PMT) detectors, three hybrid (HyD) detectors, and the Leica Application Suite AF 3.2.1.9702.

#### Physiological tests with Nitrospira enrichment cultures

To determine the optimal pH conditions for growth, three of the initial *Nitrospira* enrichments (EN_A, EN_B, EN_C) were cultured for five weeks with repeated nitrite additions in the mineral medium as described above. The pH was adjusted to 7.6 or 9.0 by titration with 1M HCl, monitored throughout the duration of the incubation and adjusted when necessary. All incubations were performed in duplicates. Enrichments which had been cultured at around pH 9 were incubated for an additional period of 25 days, during which the pH was raised sequentially every eight or nine days to 10, 10.5, and finally to 11 by titration with 1 M NaOH. The optimal nitrite concentration for cultivation was determined using the enrichment culture of “*Ca*. N. alkalitolerans”. The culture was incubated in the presence of 0.15, 0.3, 0.7, and 1 mM NaNO_2_ for 30 days. In all physiological experiments, formaldehyde-fixed culture aliquots were incubated as a negative control under the same conditions. The nitrite and nitrate concentrations were measured according to Miranda *et al*. [2].

#### Metagenome sequencing, Nitrospira genome assembly, and genome annotation

An Illumina sequencing library was prepared from the DNA sample of the “*Ca*. N. alkalitolerans” enrichment using the ruSeq DNA PCR-free sample preparation (Illumina) following the manufacturer’s recommendations and paired-end sequenced (2×300 bp) twice on a MiSeq using a MiSeq Reagent kit v3 (Illumina) following the manufacturer’s recommendations. Base calling was carried out using MiSeq control software v.2.5. Illumina read quality and adaptor trimming (trim limit: 0.01, no ambiguous bases, min length: 55 bp), de novo assembly (word size: 21, bubble size: 186, min length: 500 bp), and read mapping (default settings except length fraction: 0.95 and similarity fraction: 0.95) were performed in CLC Genomics Workbench v. 8.5.1. The Illumina de novo assembly was checked for contamination and completeness using the mmgenome workflow (http://madsalbertsen.github.io/mmgenome/).

The most abundant scaffolds in the assembly were identified as Nitrospirae and binned into a provisional metagenome assembled genome (MAG) using mmgenome as described above. This MAG was further separated into three sub-bins based on coverage. Each of these sub-bins underwent iterative reassembly using MAGspinner (https://github.com/hexaquo/MAGspinner) which is a recently updated and automated version (used in [11]) of the manual method used in [12–14]. MAGspinner calculates null models for tetranucleotide composition and coverage for a MAG, rejects scaffolds that are outside the model and then maps (BBMap v. 36.32) raw reads to scaffolds for reassembly in Spades v. 3.10.1 [15], using the scaffolds that fit the null model as “trusted contigs”. The reassembly was repeated (in blocks of ten rounds) until genome statistics became self-consistent (based on output from CheckM v. 1.0.7 “checkm qa --tab_table -o 2”, [16]). Self-consistency was established by calculating the correlation between all checkM-calculated statistics and the number of rounds over the previous ten rounds of reassembly. When no statistic was correlated with round number (p>0.05 for all statistics), the iterative procedure ceased. Of the three *Nitrospirae* MAG sub-bins, only the most abundant remained consistently high in coverage throughout the binning procedure. The coverage of the two less abundant sub-bins “drifted” upwards over the reassembly procedure, indicating that the organisms represented by them were closely related to the organism represented by the most abundant sub-bin and that these sub-bins could not be delineated confidently. The novelty of original and reassembled sub-bins were assessed against published *Nitrospirae* using genomic average nucleotide identity (gANI) calculated with gANI-MiSI [17] and average amino acid identity (AAI) as calculated using bidirectional best blastp hits aligned over at least 70% of the length of both genes. The % identity was weighted by query gene length. The calculations were repeated using each genome as query and target and the average of both calculations was reported.

The more abundant sub-bin was accepted as the final “*Ca*. N. alkalitolerans” genome bin and uploaded to the MicroScope platform [18] under the name “*Candidatus* Nitrospira alkalitolerans strain KS” for automatic gene prediction and annotation, which was amended manually where necessary for key genes of chemolithoautotrophic nitrite oxidation, energy metabolism, and adaptations to the haloalkaline environment. Genome statistics in Table S1 were collected with the integrated tools of the MicroScope platform.

#### Identification and phylogenetic analysis of “Candidatus Nitrospira alkalitolerans” gene products putatively involved in haloalkalitolerance

Homology searches for orthologous genes were performed using the OrthoFinder software with default settings [19]. Candidate genes for pH homeostasis and osmoregulation at alkaline conditions were searched manually by browsing the automated gene annotations and by using the BLAST tool of the MicroScope platform. Amino acid alignments of ATPase subunit c genes were constructed with Clustal Omega [20] and visualized with ESPript3 [21].

#### Cloning, DNA extraction, amplicon sequencing, and phylogenetic analysis of 16S rRNA gene and nxrB amplicons

PCR products of 16S rRNA gene fragments from *Nitrospira* enrichment cultures were generated using primers 8F (5’ AGA GTT TGA TYM TGG CTC 3’) [22] and the *Nitrospira*-specific primer R1158 (5’CCC GTT MTC CTG GGC AGT 3’) [6] as described in Herbold *et al*. (2015). The 20 µl PCR reaction consisted of 1× Taq buffer (Fermentas), 0.2 mM dNTPmix (Fermentas), 2 mM MgCl_2_ (Fermentas), 0.025 U Taq DNA polymerase (Fermentas), 0.1 mg mL^−1^ bovine serum albumin, 1 μM of each of the forward and reverse primers, and 1 μL of enrichment culture that had been freeze-thawed three times. The primary melting step of the PCR assay was extended to 15 min, but otherwise cycling was conducted as performed by Herbold *et al*. [23]. The amplified, 550 bp-long Nitrospira 16S rRNA gene fragments were purified using QIAquick PCR Purification Kit (Qiagen) and cloned with the TOPO-TA cloning Kit (Fisher Scientific) according to the manufacturers’ instructions. Plasmids from successfully transformed *E. coli* cells were harvested with the Plasmid MiniPrep Kit (Quiagen) according to the manufacturer’s instructions. The cloned inserts were PCR-amplified using the M13 forward and reverse primers (included in the TOPO-TA cloning kit) according to the manufacturer’s instructions and were then Sanger sequenced (Microsynth Austria GmbH). Enrichments were screened for the presence of Nitrospina by PCR targeting the nxrB with primers F169/ Rb638 (5’ TAC ATG TGG TGG AAC A 3’/ 5’CGR GAC TGA TCG ATC A 3’, [17]; reverse primer: personal communication from Sebastian Lücker). PCR and cycling procedures were as described in Herbold *et al*. [23] with the exception of an annealing temperature of 55°C.

For amplicon sequencing, DNA was extracted from environmental samples according to Angel *et al*. [24] and used as template for PCR with primer 8F/R1158 (see above) targeting the 16S rRNA gene, and primers Fa169/Ra638 (see above/5’ CGG TTC TGG TCR ATC A 3’; [25]) targeting the *nxrB* of *Nitrospira*. PCR and amplicon library preparation for Illumina MiSeq sequencing were performed as described in Herbold et al. [23]. MiSeq sequencing was performed at Microsynth Austria GmbH. Paired end reads were processed and mapped to operational taxonomic unit (OTU) representatives as described in Herbold *et al*. [16] including chimera checks using the UPARSE pipeline [26, 27]. The 16S rRNA gene amplicons were 550 bp long and underwent quality filtering and trimming using a strategy that is based on the protocol for Illumina data in the Earth Microbiome Project (Version 5 2012, [28]. Resulting library sizes ranged between 1172 and 8165 obtained sequences for the 16S rRNA gene data set and between 2507 and 10613 obtained sequences for the *nxrB* data set.

The species level thresholds for OTU clustering were 97% sequence identity for the 16S rRNA gene amplicons and 95% sequence identity for *nxrB* amplicons (the latter according to Pester *et al. [25]*). Non-target sequences were identified by aligning representative sequences from each OTU to the SILVA Ref data set Nr 99, release 132 [29] or to the *nxrB* reference data set from Pester and colleagues [25] with the integrated aligner and manual curation in ARB [30] and excluded from further analysis.

A maximum likelihood tree was generated in RaxML [31] using 30 full-length 16S rRNA gene sequences of selected reference Nitrospira species that had been aligned with the integrated aligner in ARB [30]. *Leptospirillum ferrooxidans* (AJ237903), *Ca*. Magnetobacterium bavaricum (FP929063), *Thermodesulfovibrio yellowstonii* DSM 11347 (CP001147) and *Ca*. Methylomirabilis oxyfera (FP565575) were used as outgroup. A 50% conservation filter was applied, resulting in 1310 valid alignment positions. The gamma model of rate heterogeneity and the generalized time-reversible (GTR) substitution model were utilized and 100 rapid bootstrap inferences were executed. The representative 16S rRNA gene sequences from each *Nitrospira* OTU and the 16S rRNA gene sequences from the three most abundant clones, were aligned to the set of full-length sequences and added to the maximum likelihood tree with the Evolutionary Placement Algorithm (EPA) using the GTRGAMMA substitution and rate heterogeneity model on 745 alignment positions.

#### Geographical maps and Statistical analyses

Generation of geographical maps and statistical analyses and were performed using the software R [32] version 3.3.2. Maps were drawn based on the data available in the package ‘maps’ and subsequently modified with Inkscape (www.inkscape.org).

16S rRNA gene sequence libraries were normalized with the GMPR method [33] prior to further analysis. Estimates of *Nitrospira* 16S rRNA gene OTU richness and the inverse Simpson’s index as a proxy for community diversity were calculated with the functions ‘estimatR’ and ‘diversity’ of the R package vegan [34]. Environmental factors determined in this study were centered and/or scaled using the function ‘scale’ and checked for auto-correlation with the function ‘pairs’. Since the dissolved organic matter concentrations correlated with the concentrations of total dissolved nitrogen, nitrite and iron, organic matter concentrations were excluded from further analysis. *Nitrospira* communities were grouped by hierarchical clustering of a Bray-Curtis dissimilarity matrix using the Ward agglomeration method with the command ‘hclust’. Principal coordinate analysis (PCoA) of *Nitrospira* communities was performed using the ‘cmdscale’ command. Differences in environmental conditions determined in this study between lakes were assessed by Welch’s unequal variances t-Test on scaled and centered data. Significance of differences in nitrite utilization and nitrate production by alkalitolerant *Nitrospira* enrichments over time were determined by correlation analysis with the function ‘cor.test’ using the Pearson method. Significant correlations (p ≤ 0.05) between environmental factors determined in this study and *Nitrospira* OTU richness and diversity were determined with the command ‘cor.test’ using the Spearman method.

### Supplemental Results

#### Adaptations to low iron availability

Bioavailable iron is scarce in marine and alkaline systems [35, 36], and organisms living at elevated pH conditions use various mechanisms for iron sequestration [37, 38]. Like other *Nitrospira*, “*Ca*. N. alkalitolerans” must have a high demand for iron as key metabolic pathways of Nitrospira, such as nitrite oxidation, the electron transport chain, and CO_2_ fixation, depend on enzymes with iron-sulfur clusters or heme as cofactors [39]. Accordingly, the genome of “*Ca*. N. alkalitolerans” contains numerous genes for iron uptake and storage. Among them are six genes of putative TonB-dependent iron siderophore receptors, four gene copies of the energy-transducing inner membrane TonB/ExbB/ExbD complex, and several genes coding for bacterioferritin (Table S2). One of the TonB-dependent iron siderophore receptors is most similar to a homolog of the marine *Methylocaldum marinum* (47.3% AA identity). Interestingly, the genome of “*Ca*. N. alkalitolerans” does not appear to encode any known pathway for the synthesis of siderophores. Thus, we assume that this nitrite oxidizer scavenges siderophores produced by other organisms while saving the costs of siderophore biosynthesis from fixed inorganic carbon similar to *Nitrosomonas europaea* [40].

#### Adaptation to toxic arsenite

High concentrations of inorganic arsenic have been found in several soda lakes and pristine aquifers [41–44], and the inhabiting microorganisms must possess mechanisms to deal with the presence of this highly toxic metal. The saline-alkaline lakes of the national park “Neusiedler See - Seewinkel” might also contain elevated levels of arsenite, as this has been reported for wetlands in the same area [45]. In the genome of “*Ca*. N. alkalitolerans”, we identified the gene *arsB*, which codes for a PMF-dependent arsenite resistance efflux pump of the ACR3 family [46] (Fig. 5, Table S2). Since this type of arsenite efflux pump is not present in any other characterized NOB, and the gene of “*Ca*. N. alkalitolerans” exhibits a high similarity to a homologous gene in *Nitrincola nitratireducens* (66.5% AA identity) isolated from a haloalkaline lake, its presence in the genome may be a specific adaptation of “*Ca*. N. alkalitolerans” to its extreme habitat.

