## Supplemental table S1 for "Exploring the upper pH limits of nitrite oxidation: diversity, ecophysiology, and adaptive traits of haloalkalitolerant *Nitrospira*"

**Table S1** *Nitrospira* MAG statistics

| **Attribute** | ***Ca.* Nitrospira alkalitolerans** | ***Nitrospira* sub-bin1*** | ***Nitrospira* sub-bin2*** |
|---|---|---|---|
| Size (bp) | 4.894.714 | 957.413 | 658.454 |
| Completeness (%) | 95.83 | 21.93 | 0 |
| Contamination (%) | 4.83 | 0 | 0 |
| DNA G + C (%) | 51.4 | 51 | 51.5 |
| DNA scaffolds | 87 | 293 | 217 |
| Total genes | 5107 | 1142 | 799 |
| RNA genes | 35 | 3 | 90 |
| rRNA genes | 3 | 0 | 0 |
| tRNA genes | 47 | 8 | 8 |
| Amino acids encoded by tRNA genes (unique codons) | 20 (46) | 6 (8) | 5 (8) |
| *nxrA genes (incl. Fragments)* | 3 | 4 | 3 |
| *nxrB genes (incl. Fragments)* | 1 | *1* | *0* |
| Pairwise gANI value to “Ca. N. alkalitolerans” bin | 100 | 97.1 | 95.2 |
| Oneway gAAI value to “Ca. N. alkalitolerans” bin | 99.9 | 94.3 | 81.1 |

*original sub-bin, not re-assembled
