## Supplemental table S2 for "Exploring the upper pH limits of nitrite oxidation: diversity, ecophysiology, and adaptive traits of haloalkalitolerant *Nitrospira*"

Selected proteins of “*Ca.* Nitrospira alkalitolerans” with predicted functions in metabolic pathways and in adaptation to high alkalinity and salinity

| **Central Carbon Metabolism** | | | |
|---|---|---|---|
| **Glycogen formation/degradation** | **MaGe Identifier** | **Gene** | **Product** |
| | NSPALKALI_v1_40047 | *glgX* | Glycogen debranching enzyme |
| | NSPALKALI_v1_60125 | *glgX* | Glycogen debranching enzyme |
| | NSPALKALI_v1_170014 | *glgX* | Putative Glycogen debranching enzyme |
| | NSPALKALI_v1_40023 | *glgP* | Glycogen phosphorylase |
| | NSPALKALI_v1_140097 | *glgP* | Glycogen phosphorylase |
| | NSPALKALI_v1_60127 | *glgB* | 1,4-alpha-glucan branching enzyme |
| | NSPALKALI_v1_60129 | *glgC* | Glucose-1-phosphate adenylyltransferase |
| | NSPALKALI_v1_240060 | *glgC* | Glucose-1-phosphate adenylyltransferase |
| | NSPALKALI_v1_60051 | *glgE* | Alpha-1,4-glucan:maltose-1-phosphate maltosyltransferase |
| | NSPALKALI_v1_680007 | *pgm* | Phosphoglucomutase |
| | NSPALKALI_v1_30082 | *glk* | Glucokinase |
| | NSPALKALI_v1_120061 | *pgi* | Glucosephosphate isomerase |
| | NSPALKALI_v1_40085 | *malQ* | 4-alpha-glucanotransferase |
| | NSPALKALI_v1_360024 | *rfbF* | Glucose-1-phosphate cytidylyltransferase |
| | NSPALKALI_v1_80165 | | Amylo-alpha-1,6-glucosidase |
| **(r)TCA cycle** | **MaGe Identifier** | **Gene** | **Product** |
| | NSPALKALI_v1_20059 | *aclB* | ATP-citrate lyase, beta subunit |
| | NSPALKALI_v1_20060 | *aclA* | ATP citrate lyase, alpha subunit |
| | NSPALKALI_v1_200042 | *forB* | 2-oxoglutarate:ferredoxin oxidoreductase, beta subunit |
| | NSPALKALI_v1_200043 | *forC* | 2-oxoglutarate:ferredoxin oxidoreductase, gamma subunit |
| | NSPALKALI_v1_200044 | *forE* | 2-oxoglutarate:ferredoxin oxidoreductase, epsilon subunit |
| | NSPALKALI_v1_200045 | *forD* | 2-oxoglutarate:ferredoxin oxidoreductase, delta subunit |
| | NSPALKALI_v1_200046 | *forA* | 2-oxoglutarate:ferredoxin oxidoreductase, alpha subunit |
| | NSPALKALI_v1_20019 | *porD* | Pyruvate:ferredoxin oxidoreductase, delta subunit |
| | NSPALKALI_v1_20018 | *porC* | Pyruvate:ferredoxin oxidoreductase, gamma subunit |
| | NSPALKALI_v1_20017 | *porB* | Pyruvate:ferredoxin oxidoreductase, beta subunit |
| | NSPALKALI_v1_20016 | *porA* | Pyruvate:ferredoxin oxidoreductase, alpha subunit |
| | NSPALKALI_v1_20015 | | CO dehydrogenase or Pyruvate:ferredoxin oxidoreductase, delta subunit |

| | | | |
|---|---|---|---|
| | NSPALKALI_v1_510019 | *sdhA/ nadB* | Succinate dehydrogenase/fumarate reductase, flavoprotein subunit or L-aspartate oxidase |
| | NSPALKALI_v1_20130 | *sdhA/ nadB* | Succinate dehydrogenase/fumarate reductase, flavoprotein subunit or L-aspartate oxidase |
| | NSPALKALI_v1_200063 | *sdhA/ nadB* | Succinate dehydrogenase/fumarate reductase, flavoprotein subunit or L-aspartate oxidase |
| | NSPALKALI_v1_510021 | *sdhC* | Succinate dehydrogenase, subunit C (fragment) |
| | NSPALKALI_v1_510020 | *sdhB* | Succinate dehydrogenase/ fumarate reductase iron-sulfur protein |
| | NSPALKALI_v1_510018 | *gltA* | Citrate synthase |
| | NSPALKALI_v1_750005 | *gltA* | Citrate synthase 2 |
| | NSPALKALI_v1_170072 | *sucA* | Pyruvate/2-oxoglutarate dehydrogenase complex,dehydrogenase (E1) component, eukaryotic type, alpha subunit |
| | NSPALKALI_v1_190028 | | Pyruvate/2-oxoglutarate dehydrogenase complex, dehydrogenase (E1) component, eukaryotic type, alpha subunit |
| | NSPALKALI_v1_170073 | *sucB* | Pyruvate/2-oxoglutarate dehydrogenase complex,dehydrogenase (E1) component, eukaryotic type, beta subunit |
| | NSPALKALI_v1_20057 | *icd* | Fragment of isocitrate dehydrogenase |
| | NSPALKALI_v1_40102 | *icd* | Putative isocitrate dehydrogenase (NADP) |
| | NSPALKALI_v1_40103 | *icd* | Isocitrate dehydrogenase (NAD+) |
| | NSPALKALI_v1_100028 | *fumC* | Fumarate hydratase (fumarase C),aerobic Class II |
| | NSPALKALI_v1_580009 | *mdh* | Malate dehydrogenase |
| | NSPALKALI_v1_200065 | *sucC* | Succinyl-CoA ligase [ADP-forming], subunit alpha |
| | NSPALKALI_v1_200064 | *sucD* | Succinyl-CoA ligase [ADP-forming] subunit beta |
| | NSPALKALI_v1_200061 | *acnA* | aconitate hydratase 1 |
| | NSPALKALI_v1_90082 | *lpdA* | 2-oxo-acid dehydrogenase complex, Dihydrolipoamide dehydrogenase (E3) (fragment) |
| **Carbonate uptake** | **MaGe Identifier** | **Gene** | **Product** |
| | NSPALKALI_v1_30038 | *bicA* | Bicarbonate transporter BicA or sulfate permease |
| | NSPALKALI_v1_40075 | *bicA* | Bicarbonate transporter BicA or sulfate permease |
| | NSPALKALI_v1_120088 | | Carbonic anhydase |
| | NSPALKALI_v1_550019 | | Carbonic anhydase |
| **Hydrogen Metabolism** | | | |
| **Hydrogen Metabolism** | **MaGe Identifier** | **Gene** | **Product** |
| | NSPALKALI_v1_490036 | | Putative (NiFe) hydrogenase, beta subunit (group 3b) |
| | NSPALKALI_v1_160002 | | Nickel-dependent hydrogenase, large subunit (group 3b) |
| | NSPALKALI_v1_500001 | | Hydrogenase transcriptional regulatory protein HoxA |
| | NSPALKALI_v1_500003 | | Ni,Fe-hydrogenase, small subunit (group 2a) |
| | NSPALKALI_v1_500004 | | Ni,Fe-hydrogenase, large subunit (group 2a) |
| | NSPALKALI_v1_500007 | | Hydrogenase maturation protein (fragment) |
| | NSPALKALI_v1_500014 | *hypF* | Hydrogenase maturation protein |
| | NSPALKALI_v1_500015 | *hypC* | Hydrogenase assembly chaperone protein |

| | | | |
|---|---|---|---|
| | NSPALKALI_v1_500017 | *hypD* | hydrogenase expression/ formation protein |
| | NSPALKALI_v1_500018 | *hypE* | hydrogenase expression/ formation protein |
| | NSPALKALI_v1_500020 | *hypA* | putative hydrogenase nickel incorporation protein |
| | NSPALKALI_v1_500021 | *hypB* | hydrogenase accessory protein |
| **Nitrite Metabolism** | | | |
| **Nitrite oxidation** | **MaGe Identifier** | **Gene** | **Product** |
| | NSPALKALI_v1_140038 | *nxrB* | Nitrite oxidoreductase beta subunit |
| | NSPALKALI_v1_140037 | *nxrA* | Nitrite oxidoreductase alpha subunit |
| | NSPALKALI_v1_530013 | *nxrA* | Nitrite oxidoreductase alpha subunit |
| | NSPALKALI_v1_530020 | *nxrA* | Nitrite oxidoreductase alpha subunit (fragment) |
| | NSPALKALI_v1_150108 | *nxrC* | Putative nitrite oxidoreductase, gamma subunit |
| | NSPALKALI_v1_70126 | *nxrC* | Nitrite oxidoreductase gamma subunit |
| | NSPALKALI_v1_70134 | *nxrC* | Nitrite oxidoreductase gamma subunit |
| **Respiratory Chain** | | | |
| **Complex I** | **MaGe Identifier** | **Gene** | **Product** |
| | NSPALKALI_v1_400013 | *nuoA* | NADH-quinone oxidoreductase subunit A |
| | NSPALKALI_v1_400014 | *nuoB* | NADH-quinone oxidoreductase subunit B |
| | NSPALKALI_v1_400015 | *nuoC/D* | NADH-quinone oxidoreductase subunit C/D |
| | NSPALKALI_v1_400016 | *nuoE* | NADH-quinone oxidoreductase subunit E |
| | NSPALKALI_v1_400017 | *nuoF* | NADH-quinone oxidoreductase, subunit F |
| | NSPALKALI_v1_400018 | *nuoG* | NADH-quinone oxidoreductase, subunitG |
| | NSPALKALI_v1_400019 | *nuoH* | NADH-quinone oxidoreductase subunit H |
| | NSPALKALI_v1_400020 | *nuoI* | NADH-quinone oxidoreductase subunit I |
| | NSPALKALI_v1_400021 | *nuoJ* | NADH-quinone oxidoreductase, membrane subunit J |
| | NSPALKALI_v1_400022 | *nuoK* | NADH-quinone oxidoreductase subunit K |
| | NSPALKALI_v1_400023 | *nuoL* | NADH-quinone oxidoreductase, membrane subunit L |
| | NSPALKALI_v1_400024 | *nuoM* | NADH-quinone oxidoreductase subunit M |
| | NSPALKALI_v1_400025 | *nuoN* | NADH-quinone oxidoreductase subunit N2 |
| | NSPALKALI_v1_250014 | *nuoA* | NADH-quinone oxidoreductase subunit A |
| | NSPALKALI_v1_250015 | *nuoB* | NADH-quinone oxidoreductase subunit B |
| | NSPALKALI_v1_250016 | *nuoC* | NADH-quinone oxidoreductase subunit C |
| | NSPALKALI_v1_250017 | *nuoD* | NADH-quinone oxidoreductase subunit D |
| | NSPALKALI_v1_250018 | *nuoG* | NADH-quinone oxidoreductase, subunitG |
| | NSPALKALI_v1_250019 | *nuoI* | NADH-quinone oxidoreductase subunit I |

| | | | |
|---|---|---|---|
| | NSPALKALI_v1_250020 | *nuoJ* | NADH-quinone oxidoreductase, membrane subunit J |
| | NSPALKALI_v1_250021 | *nuoK* | NADH-quinone oxidoreductase subunit K |
| | NSPALKALI_v1_250022 | *nuoL* | NADH-quinone oxidoreductase, membrane subunit L |
| | NSPALKALI_v1_250023 | *nuoM* | NADH-quinone oxidoreductase, membrane subunit M |
| | NSPALKALI_v1_250024 | *nuoM* | NADH-quinone oxidoreductase, membrane subunit M |
| | NSPALKALI_v1_250025 | *nuoN* | NADH-quinone oxidoreductase subunit N |
| | NSPALKALI_v1_580010 | *nuoF* | NADH-quinone oxidoreductase subunit F 2 |
| **Complex II** | **MaGe Identifier** | **Gene** | **Product** |
| | NSPALKALI_v1_510019 | *sdhA/ nadB* | Succinate dehydrogenase/fumarate reductase, flavoprotein subunit or L-aspartate oxidase |
| | NSPALKALI_v1_510020 | *sdhB* | Succinate dehydrogenase/ fumarate reductase iron-sulfur protein |
| | NSPALKALI_v1_510021 | *sdhC* | Succinate dehydrogenase, subunit C (fragment) |
| | NSPALKALI_v1_20130 | *sdhA/ nadB* | Succinate dehydrogenase/fumarate reductase, flavoprotein subunit or L-aspartate oxidase |
| | NSPALKALI_v1_200063 | *sdhA/ nadB* | Succinate dehydrogenase/fumarate reductase, flavoprotein subunit or L-aspartate oxidase |
| **Complex III** | **MaGe Identifier** | **Gene** | **Product** |
| | NSPALKALI_v1_150103 | | Putative Quinol-cytochrome c reductase, cytochrome b subunit |
| | NSPALKALI_v1_150102 | | Putative Quinol-cytochrome c reductase, iron-sulfur subunit, modulated with PRC-barrel (Modular protein) |
| | NSPALKALI_v1_370011 | | Menaquinol-cytochrome c reductase cytochrome b subunit |
| | NSPALKALI_v1_370010 | | Putative Quinol-cytochrome c reductase, iron-sulfur subunit (Rieske iron-sulfur protein) |
| **Complex IV** | **MaGe Identifier** | **Gene** | **Product** |
| | NSPALKALI_v1_150105 | | Putative Cytochrome bd-type quinol oxidase subunit 1 |
| | NSPALKALI_v1_530011 | | Putative Cytochrome bd-type quinol oxidase, subunit 1 |
| | NSPALKALI_v1_70108 | | Cytochrome bd ubiquinol oxidase subunit I |
| | NSPALKALI_v1_220083 | | Putative Cytochrome bd ubiquinol oxidase, subunit I |
| | NSPALKALI_v1_220084 | | Putative Cytochrome bd ubiquinol oxidase, subunit II |
| **Complex V** | **MaGe Identifier** | **Gene** | **Product** |
| | NSPALKALI_v1_10079 | *atpH* | ATP synthase subunit delta |
| | NSPALKALI_v1_10080 | *atpA* | F1 sector of membrane-bound ATP synthase, alpha subunit |
| | NSPALKALI_v1_10081 | *atpD* | Membrane-bound ATP synthase , F1 sector, beta-subunit |
| | NSPALKALI_v1_10082 | *atpC* | ATP synthase epsilon chain |
| | NSPALKALI_v1_790004 | *atpA* | ATP synthase subunit alpha 2 |
| | NSPALKALI_v1_20126 | *atpI* | Putative ATP synthase F0, subunit I |
| | NSPALKALI_v1_20125 | *atpB* | ATP synthase subunit a |
| | NSPALKALI_v1_20124 | *atpE* | ATP synthase subunit c |
| | NSPALKALI_v1_20123 | *atpF* | ATP synthase subunit b |

| | | | |
|---|---|---|---|
| | NSPALKALI_v1_120045 | *atpG* | ATP synthase gamma chain |
| | NSPALKALI_v1_330007 | *atpG* | ATP synthase gamma chain |
| | NSPALKALI_v1_10070 | *yidC* | Membrane protein insertase YidC |
| **Iron uptake and storage** | | | |
| | **MaGe Identifier** | **Gene** | **Product** |
| | NSPALKALI_v1_170059 | | Bacterioferritin (modular protein) |
| | NSPALKALI_v1_60072 | | Bacterioferritin, iron storage and detoxification protein |
| | NSPALKALI_v1_80012 | *fecE* | Iron-dicitrate transporter |
| | NSPALKALI_v1_110032 | *fur* | Ferric uptake regulation protein |
| | NSPALKALI_v1_10210 | *fur* | Ferric uptake regulation protein |
| | NSPALKALI_v1_58005 | | putative Bacterioferritin-associated ferredoxin |
| | NSPALKALI_v1_460022 | | putative TonB-dependent receptor |
| | NSPALKALI_v1_250054 | | putative TonB-dependent receptor |
| | NSPALKALI_v1_570021 | | putative TonB-dependent receptor |
| | NSPALKALI_v1_110038 | | putative TonB-dependent receptor |
| | NSPALKALI_v1_340011 | | putative TonB-dependent receptor |
| | NSPALKALI_v1_400004 | | putative Protein tonB2 |
| | NSPALKALI_v1_400007 | *exbB* | Biopolymer transport protein |
| | NSPALKALI_v1_570025 | *exbD* | Biopolymer transport protein |
| | NSPALKALI_v1_570024 | *exbD* | Biopolymer transport protein |
| | NSPALKALI_v1_140114 | *exbD* | Biopolymer transport protein |
| **Flagellum biosynthesis** | | | |
| **Flagellum assembly** | **MaGe Identifier** | **Gene** | **Product** |
| | NSPALKALI_v1_190007 | *motA* | Flagellar motor protein MotA |
| | NSPALKALI_v1_190008 | *motB* | Flagellar motor protein MotB |
| | NSPALKALI_v1_140115 | | MotA/TolQ/ExbB proton channel family protein |
| | NSPALKALI_v1_30036 | *ompA/ motB* | putative OmpA/MotB |
| **$K^+$ transport** | | | |
| **$K^+$ uptake** | **MaGe Identifier** | **Gene** | **Product** |
| | NSPALKALI_v1_150036 | | putative potassium channel NAD-binding component |
| | NSPALKALI_v1_270047 | *trkC* | Trk system potassium uptake protein C |
| | NSPALKALI_v1_170093 | *trkB* | Trk system potassium uptake protein B |
| | NSPALKALI_v1_100131 | *trkB* | Trk system potassium uptake protein B |
| **$K^+$ efflux** | **MaGe Identifier** | **Gene** | **Product** |

| | | | |
|---|---|---|---|
| | NSPALKALI_v1_160038 | *kefB* | putative K(+) efflux antiporter KefB |
| | NSPALKALI_v1_30114 | *kefB* | Putative Glutathione-regulated potassium-efflux system |
| | NSPALKALI_v1_30115 | *kefB* | Putative Glutathione-regulated potassium-efflux system |
| **$Na^+$ transport** | | | |
| **Sodium import** | **MaGe Identifier** | **Gene** | **Product** |
| | NSPALKALI_v1_50050 | | Sodium/solute symporter, putative (fragment) |
| | NSPALKALI_v1_140059 | | Sodium:dicarboxylate symporter |
| | NSPALKALI_v1_150072 | | Uncharacterized sodium-dependent transporter YhdH |
| | NSPALKALI_v1_110138 | *agcS* | Sodium/alanine symporter |
| | NSPALKALI_v1_30038 | *bicA* | Bicarbonate transporter BicA or sulfate permease |
| | NSPALKALI_v1_40075 | *bicA* | Bicarbonate transporter BicA or sulfate permease |
| **Sodium/ cation exchange** | **MaGe Identifier** | **Gene** | **Product** |
| | NSPALKALI_v1_380014 | | Sodium/calcium exchanger |
| | NSPALKALI_v1_90107 | *nhaA* | sodium:proton antiporter |
| | NSPALKALI_v1_30009 | *nhaB* | fragment of sodium:proton antiporter (part 2) |
| | NSPALKALI_v1_30007 | *nhaB* | fragment of sodium:proton antiporter (part 1) |
| | NSPALKALI_v1_30027 | *mrpE* | putative Monovalent cation/H+ antiporter, subunit E |
| | NSPALKALI_v1_30028 | *mrpF* | putative Monovalent cation/H+ antiporter, subunit F |
| | NSPALKALI_v1_30029 | *mrpG* | putative Monovalent cation/H+ antiporter, subunit G |
| | NSPALKALI_v1_30030 | *mrpA* | putative Monovalent cation/H+ antiporter, subunit A (Fragment, part 3) |
| | NSPALKALI_v1_30031 | *mrpB* | putative Monovalent cation/H+ antiporter, subunit B |
| | NSPALKALI_v1_30032 | *mrpC* | putative Monovalent cation/H+ antiporter, subunit C |
| | NSPALKALI_v1_30033 | *mrpD* | putative Monovalent cation/H+ antiporter, subunit D |
| | NSPALKALI_v1_30034 | *mrpA/D (nuoL)* | putative Monovalent cation/H+ antiporter, subunit A or D |
| | NSPALKALI_v1_30035 | *mrpA/D (nuoM)* | putative Monovalent cation/H+ antiporter, subunit A or D |
| | NSPALKALI_v1_30037 | *nhaB* | sodium:proton antiporter |
| | NSPALKALI_v1_40037 | | Sodium/hydrogen exchanger |
| | NSPALKALI_v1_40053 | *mrpD* | putative Monovalent cation/H+ antiporter, subunit D |
| | NSPALKALI_v1_40054 | *mrpC* | putative Monovalent cation/H+ antiporter, subunit C |
| | NSPALKALI_v1_40055 | *mrpB* | putative Monovalent cation/H+ antiporter, subunit B |
| | NSPALKALI_v1_40056 | *mrpA/B* | putative Monovalent cation/H+ antiporter, subunit A or B |
| | NSPALKALI_v1_40057 | *mrpG/ phaG* | Monovalent cation/proton antiporter, MnhG/PhaG subunit |
| | NSPALKALI_v1_40058 | *mrpF* | putative Monovalent cation/H+ antiporter, subunit F (Multiple resistance and pH regulation protein F) |
| | NSPALKALI_v1_40059 | *mrpE* | putative Monovalent cation/H+ antiporter, subunit E |

| Na+ translocating complex I | | | |
|---|---|---|---|
| **Alternative complex I** | **MaGe Identifier** | **Gene** | **Product** |
| | NSPALKALI_v1_330010 | *nqrA* | Na(+)-translocating NADH-quinone reductase subunit A |
| | NSPALKALI_v1_330011 | *nqrB* | Na(+)-translocating NADH-quinone reductase subunit B |
| | NSPALKALI_v1_330013 | *nqrC* | Na(+)-translocating NADH-quinone reductase subunit C |
| | NSPALKALI_v1_330014 | *nqrD* | Na(+)-translocating NADH-quinone reductase subunit D |
| | NSPALKALI_v1_330015 | *nqrE* | Na(+)-translocating NADH-quinone reductase subunit E |
| | NSPALKALI_v1_330016 | *nqrF* | Na(+)-translocating NADH-quinone reductase subunit F |
| **cbb3-type complex IV** | | | |
| **Alternative complex IV** | **MaGe Identifier** | **Gene** | **Product** |
| | NSPALKALI_v1_40069 | | Putative Cytochrome c oxidase (cbb3-type), fused subunits I, II, and III |
| **N-type ATPase** | | | |
| **Alternative Complex V** | **MaGe Identifier** | **Gene** | **Product** |
| | NSPALKALI_v1_440034 | *atpD* | ATP synthase subunit beta |
| | NSPALKALI_v1_440035 | *atpC* | H(+)-transporting ATP synthase, subunit epsilon |
| | NSPALKALI_v1_440036 | *atpQ* | H(+)-transporting ATP synthase subunit q |
| | NSPALKALI_v1_440037 | *atpR* | F1/F0 ATPase, Methanosarcina type, subunit r |
| | NSPALKALI_v1_440038 | *atpB* | ATP synthase subunit a |
| | NSPALKALI_v1_790001 | *atpB* | ATP synthase subunit a |
| | NSPALKALI_v1_790002 | *atpE* | ATP synthase subunit c 2 |
| | NSPALKALI_v1_790003 | *atpF* | ATP synthase subunit b 2 |
| | NSPALKALI_v1_790004 | *atpA* | ATP synthase subunit alpha 2 |
| | NSPALKALI_v1_330002 | *atpG* | ATP synthase subunit gamma |
| **Osmoprotectants** | | | |
| **Glutamate synthesis** | **MaGe Identifier** | **Gene** | **Product** |
| | NSPALKALI_v1_70075 | *gltB* | Glutamate synthase [NADPH], large subunit |
| | NSPALKALI_v1_70076 | *gltD* | Glutamate synthase [NADPH], small chain |
| | NSPALKALI_v1_140012 | *glt* | Putative glutamate synthase (Zinc finger CDGSH-type domain protein) |
| | NSPALKALI_v1_40009 | *glt* | Putative glutamate synthase (Zinc finger CDGSH-type domain protein) |
| **Glycine betaine import** | **MaGe Identifier** | **Gene** | **Product** |
| | NSPALKALI_v1_140025 | *opuD* | Glycine betaine transporter |
| | NSPALKALI_v1_20246 | *opuCB* | ABC-type glycine betaine transport, periplasmic subunit |
| **Trehalose synthesis** | **MaGe Identifier** | **Gene** | **Product** |
| | NSPALKALI_v1_60049 | *treS* | Trehalose synthase |

| Other | | | |
|---|---|---|---|
| **Pathway** | **MaGe Identifier** | **Gene** | **Product** |
| Membrane adaptation | NSPALKALI_v1_490030 | *cls* | Cardiolipin synthase |
| Arsenical resistance | NSPALKALI_v1_370005 | *arsB/ acr3* | Arsenite efflux pump ArsB, ACR3 family |
