## Supplemental table S3 for "Exploring the upper pH limits of nitrite oxidation: diversity, ecophysiology, and adaptive traits of haloalkalitolerant *Nitrospira*"

**Table S3** pH monitoring and adjustments during the pH experiments with enrichment cultures A, B and C

| Day | Treatment pH | pH measured | pH adjusted to |
|---|---|---|---|
| 0 | 7.6 | 7.63 ± 0.02 | - |
| 3 | 7.6 | 7.86 ± 0.12 | 7.61 ± 0.02 |
| 5 | 7.6 | 7.84 ± 0.1 | 7.64 ± 0.04 |
| 8 | 7.6 | 7.78 ± 0.06 | 7.63 ± 0.01 |
| 11 | 7.6 | 7.82 ± 0.09 | 7.61 ± 0.02 |
| 15 | 7.6 | 7.75 ± 0.07 | 7.62 ± 0.03 |
| 17 | 7.6 | 7.78 ± 0.04 | 7.6 ± 0.02 |
| 22 | 7.6 | 7.82 ± 0.08 | 7.62 ± 0.01 |
| 27 | 7.6 | 7.79 ± 0.06 | 7.62 ± 0.01 |
| 31 | 7.6 | 7.76 ± 0.09 | 7.62 ± 0.02 |
| 35 | 7.6 | 7.78 ± 0.06 | 7.61 ± 0.03 |
| 0 | 9 | 9.03 ± 0.04 | - |
| 3 | 9 | 9.02 ± 0.06 | - |
| 8 | 9 | 9.03 ± 0.05 | - |
| 11 | 9 | 9 ± 0.06 | - |
| 17 | 9 | 9.02 ± 0.07 | - |
| 27 | 9 | 9.04 ± 0.06 | - |
| 35 | 9 | 9.02 ± 0.08 | - |
| 0 | 10 | 10 ± 0.02 | - |
| 2 | 10 | 9.97 ± 0.04 | - |
| 4 | 10 | 10 ± 0.03 | - |
| 6 | 10 | 9.99 ± 0.04 | - |
| 9 | 10 | 9.98 ± 0.02 | - |
| 9 | 10.5 | 10.52 ± 0.03 | - |
| 10 | 10.5 | 10.24 ± 0.07 | 10.51 ± 0.05 |
| 11 | 10.5 | 10.49 ± 0.05 | - |
| 14 | 10.5 | 10.36 ± 0.08 | 10.52 ± 0.02 |
| 17 | 10.5 | 10.51 ± 0.06 | - |
| 17 | 11 | 11.02 ± 0.04 | - |
| 18 | 11 | 10.72 ± 0.13 | 10.98 ± 0.03 |
| 19 | 11 | 10.97 ± 0.05 | - |
| 20 | 11 | 10.87 ± 0.09 | 11.02 ± 0.02 |
| 22 | 11 | 10.98 ± 0.03 | - |
| 25 | 11 | 10.98 ± 0.04 | - |

Values are averages (n=6) with standard deviation
