## Supplementary material for "Exploring the upper pH limits of nitrite oxidation: diversity, ecophysiology, and adaptive traits of haloalkalitolerant *Nitrospira*": Figure S1

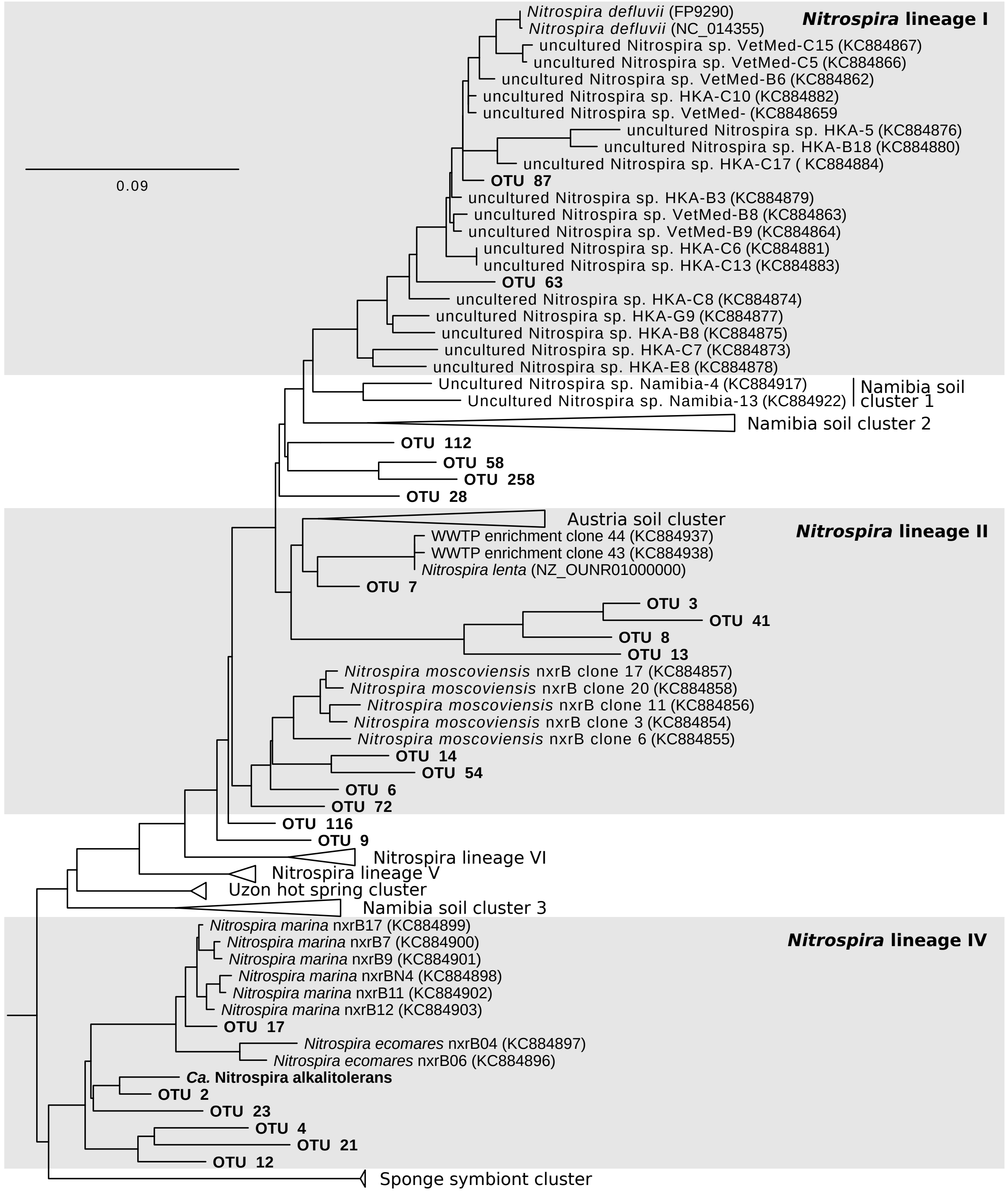


**Figure S1** Phylogenetic maximum likelihood analysis showing the affiliation of *Nitrospira nxrB* nucleotide sequences. The tree contains selected representatives from the genus *Nitrospira* and *Nitrospira* members detected in sediments of saline-alkaline lakes of the national park “Neusiedler See – Seewinkel”, Burgenland, Austria. Sequences obtained in this study are printed in bold. *Ca.* Nitrospira alkalitolerans is the organism cultured and analyzed in this study. The tree is based on the reference *nxrB* phylogeny published by Pester et al. [25]. The scale bar indicates 9% estimated sequence divergence. Please note that *Nitrospira* lineage II is usually polyphyletic in *nxrB*-based trees [25], whereas lineage II is monophyletic in 16S rRNA gene-based trees that were originally used to define the *Nitrospira* lineages I to VI [5] (see also Fig. 2 in the main text).
