## Supplementary material for "Exploring the upper pH limits of nitrite oxidation: diversity, ecophysiology, and adaptive traits of haloalkalitolerant *Nitrospira*": Figure S2

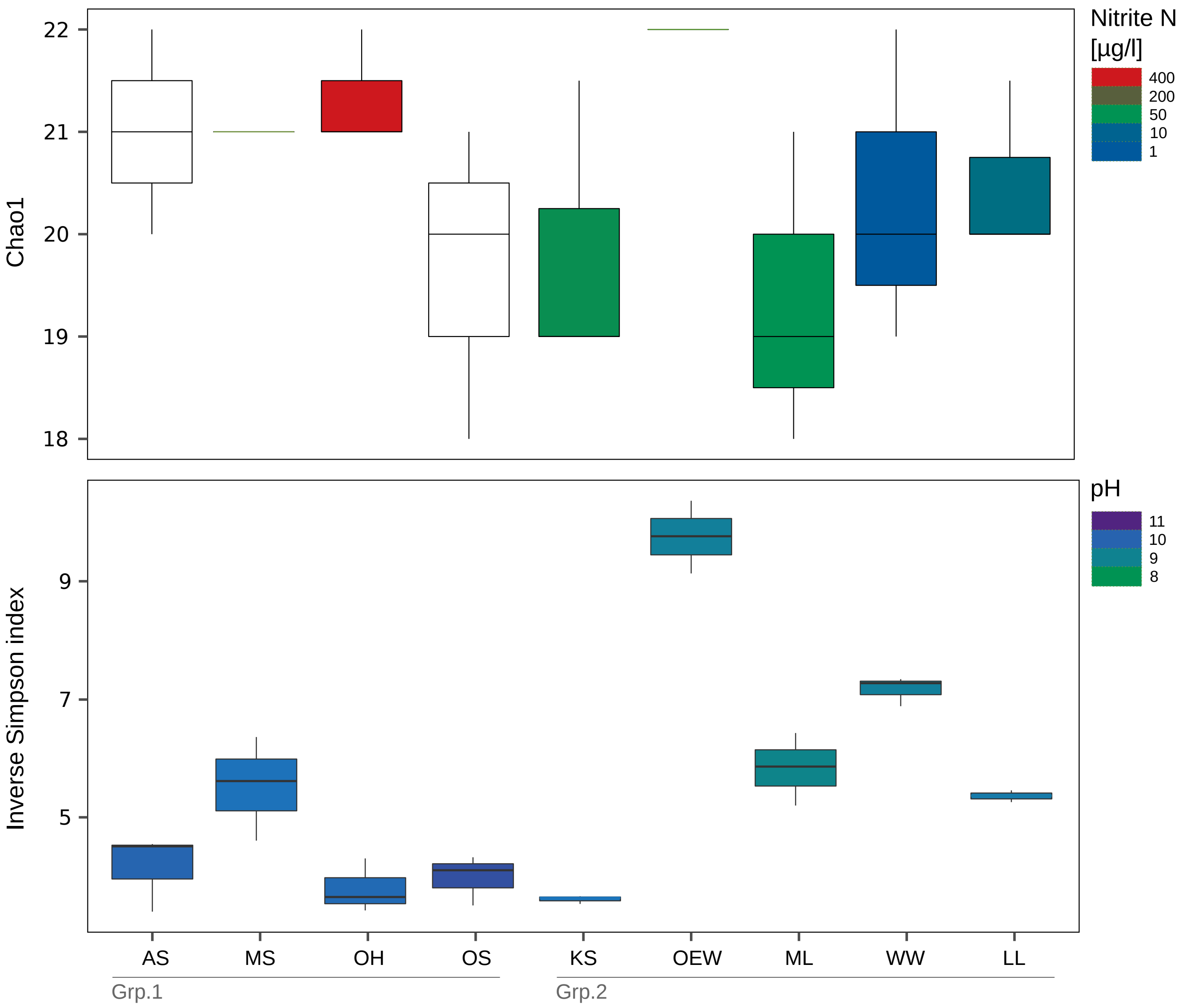


**Figure S2** Estimations of alpha diversity (richness indicator Chao1 and inverse Simpson's index) for the *Nitrospira* OTUs from the analyzed saline-alkaline lakes. The *Nitrospira* OTUs were retrieved from the 16S rRNA gene amplicon dataset using a sequence identity threshold of 97%. Boxes are colored according to nitrite concentrations and pH values in lakes in the the upper and lower panel, respectively. Non-colored boxes denote missing data. Abbreviations for the lakes are the same as in main text Table 1; Grp.1, group 1 lakes; Grp.2, group 2 lakes (see also main text Fig. 1).
