## Supplementary material for "Exploring the upper pH limits of nitrite oxidation: diversity, ecophysiology, and adaptive traits of haloalkalitolerant *Nitrospira*": Figure S3

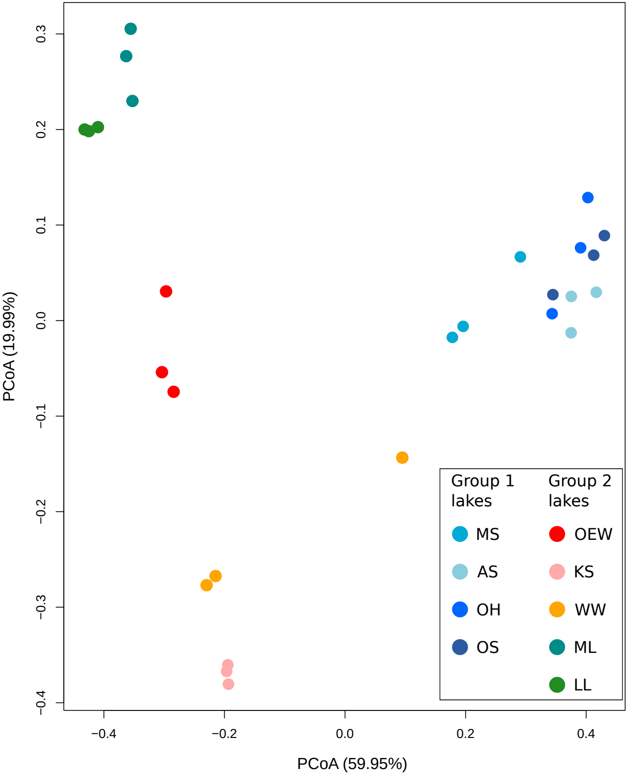


**Figure S3** Biplot showing the first two axes of a principal coordinate analysis of the *Nitrospira* communities (OTUs) in nine saline-alkaline lakes grouped by hierarchical clustering of a Bray-Curtis dissimilarity matrix using the Ward agglomeration method. The axes cumulatively explain 79.94% of the observed variation. *Nitrospira* OTUs were retrieved from the 16S rRNA gene amplicon dataset using a sequence identity threshold of 97%. Lake identifiers are the same as in Table 1 and grouping of lakes is as shown in Figure 1 of the main text.
