## Supplementary material for "Exploring the upper pH limits of nitrite oxidation: diversity, ecophysiology, and adaptive traits of haloalkalitolerant *Nitrospira*": Figure S4

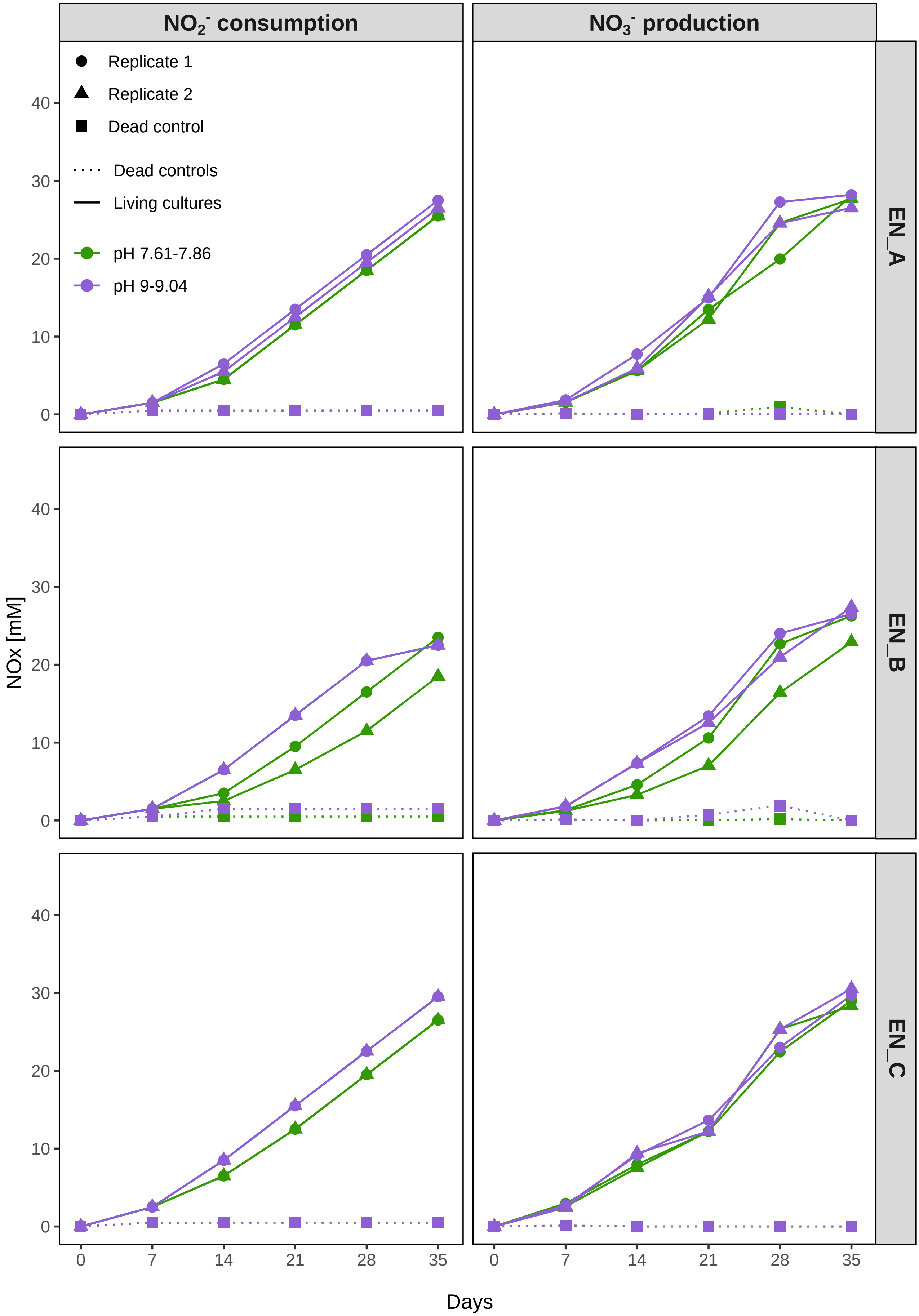


**Figure S4** Cumulative nitrite consumption and nitrate production, during 35 days by three alkalitolerant *Nitrospira* enrichment cultures from saline-alkaline lakes. The cultures were grown in mineral nitrite medium at pH 7.61-7.86 and 9-9.04. Data from two replicate incubations and one dead biomass control per pH treatment and enrichment are shown. Some symbols of replicate incubations or dead controls appear on top of each other. The pH was monitored and adjusted when necessary throughout the i ncubations (see table S3). EN_A, *Nitrospira* enrichment A; EN_B, *Nitrospira* enrichment B; EN_C, *Nitrospira* enrichment C.
