## Supplementary material for "Exploring the upper pH limits of nitrite oxidation: diversity, ecophysiology, and adaptive traits of haloalkalitolerant *Nitrospira*": Figure S5

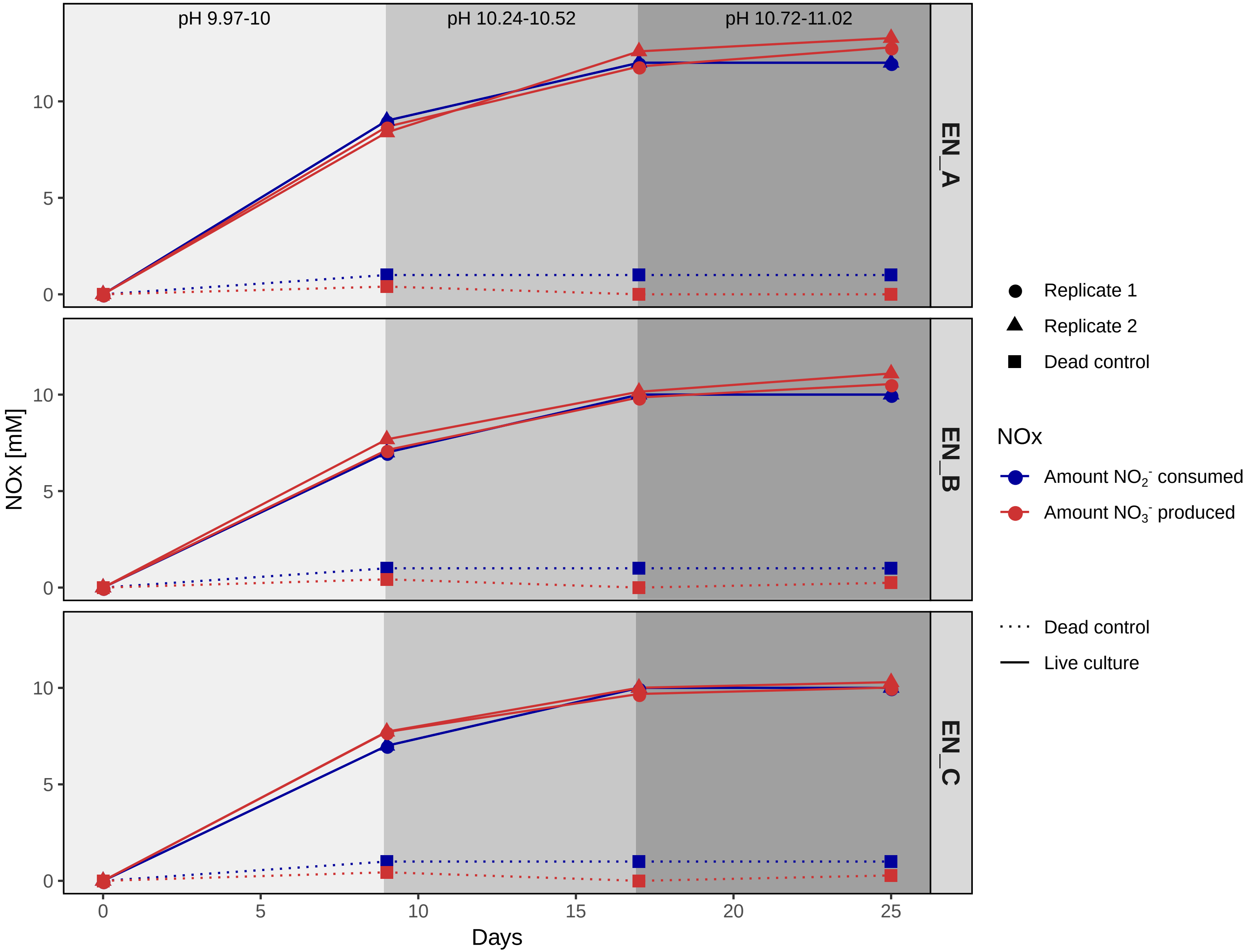


**Figure S5** Cumulative nitrite consumption and nitrate production at elevated pH conditions (up to pH 11) by three alkalitolerant *Nitrospira* enrichment cultures from saline-alkaline lakes. The cultures were grown in mineral nitrite medium and the pH was sequentially raised from 9.97-10 to 10.24-10.52 and to 10.72-11.02. The pH was monitored and adjusted when necessary throughout the incubations (see table S3). Data from two replicate incubations and one dead biomass control per pH treatment are shown. Some symbols of replicate incubations appear on top of each other. EN-A, *Nitrospira* enrichment A; EN-B, *Nitrospira* enrichment B; EN-C, *Nitrospira* enrichment C.
