## Supplementary material for "Exploring the upper pH limits of nitrite oxidation: diversity, ecophysiology, and adaptive traits of haloalkalitolerant *Nitrospira*": Figure S6

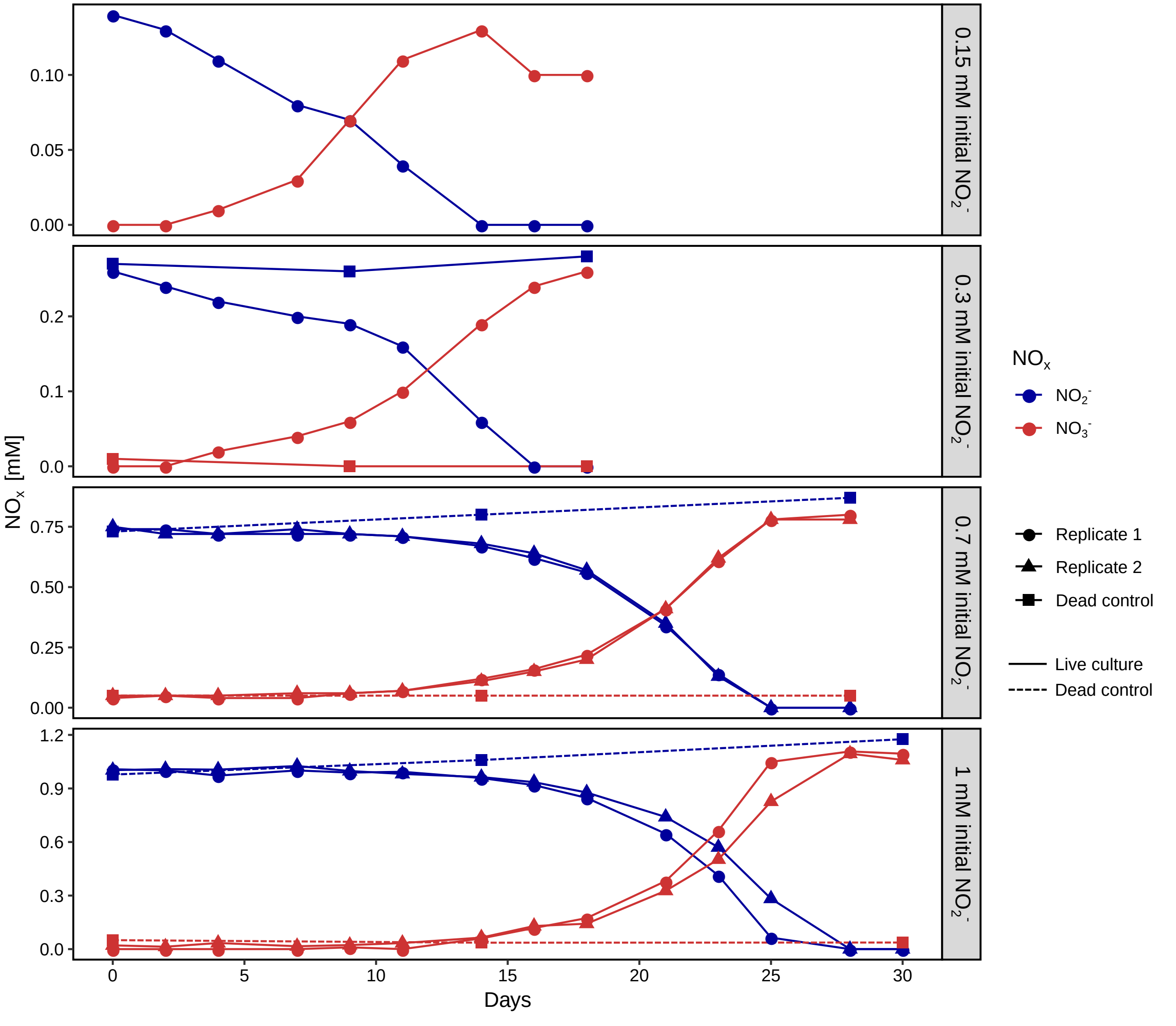


**Figure S6** Nitrite consumption and nitrate production of the "*Ca.* Nitrospira alkalitolerans" enrichment culture when cultured at pH 10.2 in mineral nitrite medium with 0.15, 0.3, 0.7, or 1 mM initial nitrite concentration. The pH was monitored and adjusted when necessary throughout the incubations. Treatments with 0.15 and 0.3 mM nitrite could not be replicated due to a lack of sufficient culture biomass.
