## Supplementary material for "Exploring the upper pH limits of nitrite oxidation: diversity, ecophysiology, and adaptive traits of haloalkalitolerant *Nitrospira*": Figure S7

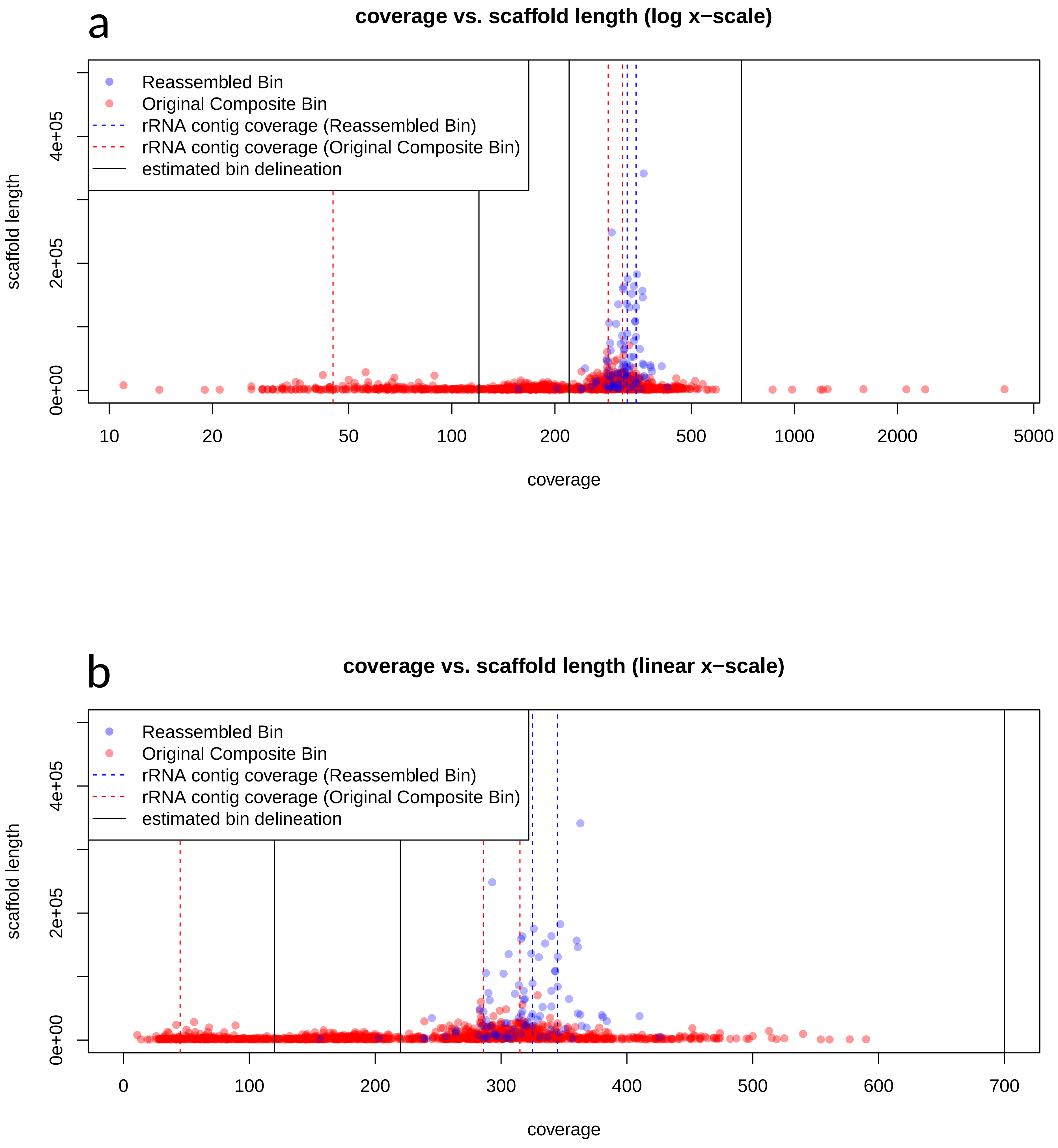


**Figure S7** Median base coverage and scaffold length of scaffolds in original *Nitrospira* MAG and the re-assembled "*Ca*. N. alkalitolerans" MAG from Illumina MiSeq sequencing of the "*Ca.* N. alkalitolerans" enrichment culture. Scaffold length (x-axis) is shown in log-scale in (a) and linear scale in (b). Contigs with coverage values of 220 – 700 were binned and re-assembled as the "*Ca.* Nitrospira alkalitolerans" MAG, while contigs with coverage values of 120 – 220 and < 120 were binned into *Nitrospira* bin1 and bin2, respectively (see also Table S1).
