## Supplementary material for "Exploring the upper pH limits of nitrite oxidation: diversity, ecophysiology, and adaptive traits of haloalkalitolerant *Nitrospira*": Figure S8

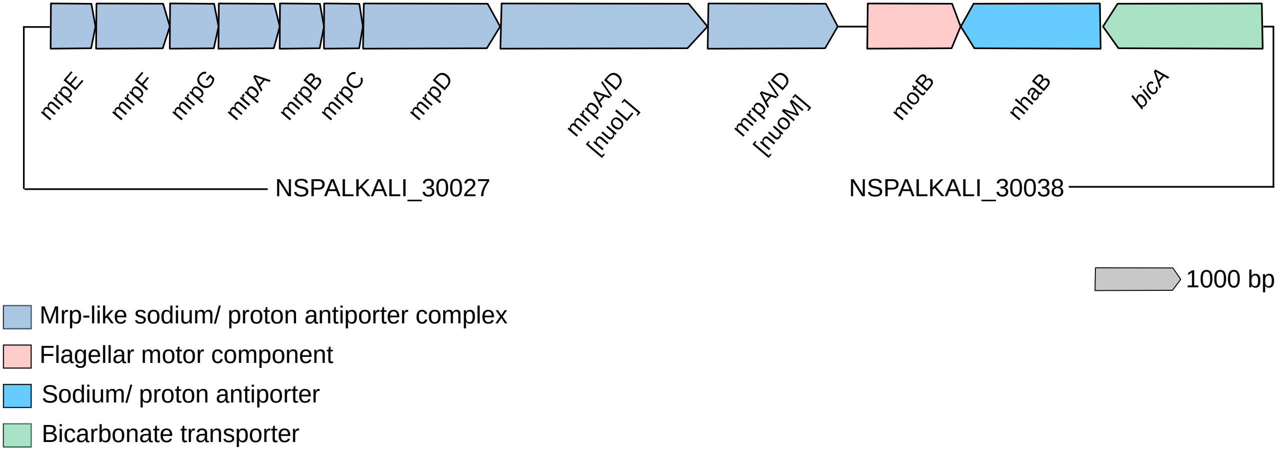


**Figure S8** Schematic illustration of one of the genomic loci with a Mrp-like sodium/proton antiporter (*mrpA-E*) in "*Ca*. N. alkalitolerans". Two additional putative MrpA or MrpD subunits display sequence similarity to the complex I subunits NuoL and NuoM, respectively. Genes are drawn to scale.
