## Supplementary material for "Exploring the upper pH limits of nitrite oxidation: diversity, ecophysiology, and adaptive traits of haloalkalitolerant *Nitrospira*": Figure S9

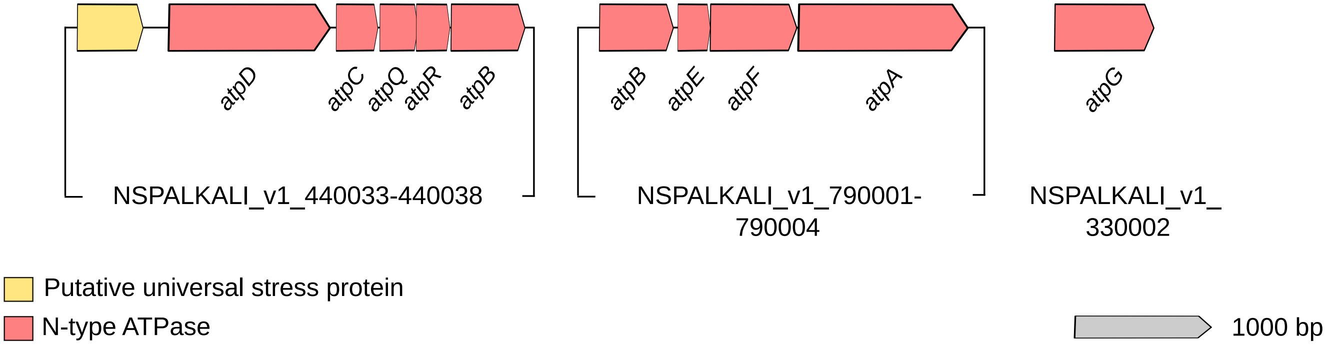


**Figure S9** Schematic illustration of the genomic loci of the N-type ATPase with genes *atpA*-*G* of "*Ca.* N. alkalitolerans". Genes are drawn to scale.
