## Supplementary material for "Exploring the upper pH limits of nitrite oxidation: diversity, ecophysiology, and adaptive traits of haloalkalitolerant *Nitrospira*": Figure S10

```
                                 HELIX N                                        LOOP
                                 1         10        20        30        40
Ca. N. alkalit.  N-type c  ...MDSMTLIAVASIVTAGITTGVGTIGPALGEGRAVSTALTSLAQQPD
B. pseu.         N-type c  .....MNNLIEVVSIAAAALAVSFGAIGPALAEGRAVGAAMDAIARQPD
M. barkeri       N-type c  MALDTYITTIAVASIATAGITIGIGVIGPAIGEGRAVATALSSLAQQPD
D. baculatum     N-type c  ...MDSMTIIAVASIIIAGITTGFGTMGPALAEGKAVATALTSLAQQPD
N. sp. Is79A3    N-type c  ...MDSMTIIAVASIITAGMTIGIGVIGPSLGEGKAVATALTSLAQQPD
F. nuc. nuc.     V-type c  MDLLTAKTIVLGCSAVGAGL.AMIAGLGPGIGEGYAAGKAVESVARQPE
I. polyt.        V-type c  MDMLLAKTVVLAASAVGAGT.AMIAGIGPGVGQGYAAGKAVESVARQPE
E. coli          F-type c  ME.....NLMNDLLYMAAAVMMGLAAIGAAIGIGILGGKFLEGAARQPD
N. def.          F-type c  MD.......AAAAALVGMGL.AAAGFAGAGVGIGYIFGKMIEAVARQPE
Ca. N. alkalit.  F-type c  MD.......SAAAALLGMGL.AAAGFAGAGIGIGYIFGKMIEAVARQPE

                                 HELIX C
                             50        60        70        80        90
Ca. N. alkalit.  N-type c  AANTITRTLFVGLAMIESTAIYCFVVSMILIFANPFWNHVLAQAAGK   H+/Na+
B. pseu.         N-type c  ASGTVSRTLFVGLAMIETMAIYCLVVALLLLFANPFVK.........   H+
M. barkeri       N-type c  ASATITRTLFVGLAMIESLAIYCFVVSMILIFANPFWNRALT.....   H+/Na+
D. baculatum     N-type c  ASATITRTLFVGLAMIESTAIYCFVVSMILIFANPFWNYAIAQMAGK   H+/Na+
N. sp. Is79A3    N-type c  ASATITRTLFVGLAMIESTAIYCFVVSMILLFANPFWNQVITQAAGK   H+/Na+
F. nuc. nuc.     V-type c  ARGSIISTMILGQAVAESTGIYSLVIALILLYANPFLSKLG......   Na+
I. polyt.        V-type c  AKGDIISTMVLGQAVAESTGIYSLVIALILLYANPFVGLLG......   Na+
E. coli          F-type c  LIPLLRTQFFIVMGLVDAIPMIAVGLGLYVMFAVA............   H+
N. def.          F-type c  AEGRVGKYMWIGFALVEAIALYGLVIAFIIMGLRK............   H+
Ca. N. alkalit.  F-type c  AEGRVGKYMWIGFALVEAIALYGLVIAFIIMGKG.............   H+
```

**Figure S10** Amino acid alignment of selected c subunits of N-, F-, and V-type ATPases including the N-and F-type c subunits identified in "*Ca.* Nitrospira alkalitolerans". The type of ATPase and the transported cations are indicated at the beginning and end of each sequence, respectively. ATPases that have not yet been specifically characterized with respect to cations transported are tagged with $H^+/Na^+$. The glutamic acid and glutamine residues in the N-terminal helix putatively serving as $Na^+$ ligands and the typical ESTxxY $Na^+$-binding motif in the C-terminal helix are highlighted in orange. Conserved regions are highlighted in grey. Included species are: "*Ca.* Nitrospira alkalitolerans" (numbering), *Burkholderia pseudomallei* 668, *Methanosarcina barkeri* Fusaro, *Desulfomicrobium baculatum* DSM 4028, *Nitrosomonas* sp. Is79A3, *Fusobacterium nucleatum* subsp. nucleatum ATCC 25586, *Ilyobacter polytropus* DSM 2926, *Escherichia coli* 042, and *Nitrospira defluvii*.
